# Immunologic and Biophysical Features of the BNT162b2 JN.1- and KP.2-Adapted COVID-19 Vaccines

**DOI:** 10.1101/2024.11.04.621927

**Authors:** Wei Chen, Kristin R. Tompkins, Ian W. Windsor, Lyndsey T. Martinez, Minah Ramos, Weiqiang Li, Shikha Shrivastava, Swati Rajput, Jeanne S. Chang, Parag Sahasrabudhe, Kimberly F. Fennell, Thomas J. McLellan, Graham M. West, Kristianne P. Dizon, Aaron Yam, Siddartha Mitra, Subrata Saha, Daiana Sharaf, Andrew P. McKeen, Carla I. Cadima, Alexander Muik, Wesley Swanson, Raquel Munoz Moreno, Pilar Mendoza Daroca, Ugur Sahin, Annaliesa S. Anderson, Huixian Wu, Kena A. Swanson, Kayvon Modjarrad

## Abstract

Vaccines remain a vital public health tool to reduce the burden of COVID-19. COVID-19 vaccines that are more closely matched to circulating SARS-CoV-2 lineages elicit more potent and relevant immune responses that translate to improved real-world vaccine effectiveness. The rise in prevalence of the Omicron JN.1 lineage, and subsequent derivative sublineages such as KP.2 and KP.3, coincided with reduced neutralizing activity and effectiveness of Omicron XBB.1.5-adapted vaccines. Here, we characterized the biophysical and immunologic attributes of BNT162b2 JN.1- and KP.2-adapted mRNA vaccine-encoded spike (S) protein immunogens. Biophysical interrogations of S revealed the structural consequences of hallmark amino acid substitutions and a potential molecular mechanism of immune escape employed by JN.1 and KP.2. The vaccine candidates were evaluated for their immunogenicity when administered as fourth or fifth doses in BNT162b2-experienced mice or as a primary series in naïve mice. In both vaccine-experienced and naïve settings, JN.1- and KP.2-adapted vaccines conferred improved neutralizing responses over the BNT162b2 XBB.1.5 vaccine against a broad panel of emerging JN.1 sublineages, including the predominant KP.3.1.1 and emerging XEC lineages. Antigenic mapping of neutralizing responses indicated greater antigenic overlap of JN.1- and KP.2-adapted vaccine responses with currently circulating sublineages compared to an XBB.1.5-adapted vaccine. CD4^+^ and CD8^+^ T cell responses were generally conserved across all three vaccines. Together, the data support the selection of JN.1- or KP.2-adapted vaccines for the 2024-25 COVID-19 vaccine formula.

**ONE-SENTENCE SUMMARY:** The Omicron JN.1- and KP.2-adapted BNT162b2 mRNA vaccines encoding prefusion S proteins elicit similar preclinical neutralizing antibody responses against circulating JN.1 sublineage pseudoviruses that are more potent than those elicited by past iterations of BNT162b2 licensed vaccines, thus demonstrating the importance of annual strain changes to the COVID-19 vaccine.

## INTRODUCTION

As coronavirus disease-2019 (COVID-19) transitions from a pandemic to an endemic state, questions remain around the evolutionary trajectory of its causative pathogen, Severe Acute Respiratory Syndrome Coronavirus-2 (SARS-CoV-2), the periodicity of its incidence, and the necessary frequency of variant-adapted vaccine updates to ensure optimal protection against a range of clinical outcomes. It remains clear, however, that the annual burden of COVID-19 cases, hospitalizations and deaths still places it among the leading global causes of infectious disease morbidity and mortality, similar to or higher than pre-pandemic levels observed for influenza and other pneumonias ^1–4^.

Since the emergence of the first SARS-CoV-2 Omicron variant of concern (VOC) in late 2021, the virus has differentiated into more than 4300 distinct genetic lineages ^5^. As these lineages have trended toward increased transmissibility and greater antigenic distance from the original ancestral strain, approved vaccines have been updated to include variant-specific mutations in the SARS-CoV-2 spike (S) ^6–10^, to maintain optimal protection against the most relevant lineages causing COVID-19. Although updated vaccines have been typically introduced to the general public during the early Autumn season of the Northern Hemisphere, vaccine manufacturers must conduct at-risk development activities throughout the preceding year in anticipation of the variant composition recommendations issued by regulators and international normative bodies in the Spring of that year ^9–11^.

During the Northern Hemisphere Winter of 2023 to 2024, the variant epidemiology of SARS-CoV-2 shifted from a dominance of Omicron XBB derivative lineages to BA.2.86 descendants, most notably the variant of interest, JN.1 ^12^ and its sublineages, achieving a peak prevalence that exceeded all prior Omicron lineages since the emergence of BA.1 and BA.2. As the global dominance of JN.1 plateaued and declined in prevalence, derivative sublineages acquired additional mutations, translating to amino acid substitutions in the S protein that conferred improved viral fitness or immune escape. Some of those sublineages (*e.g.*, KP.2) gained epidemiologic relevance during the summer months of 2024, particularly those that contained convergent F456L and R346T amino acid substitutions in the S receptor binding domain (RBD). More recently, the KP.3 sublineage, which contains the F456L, but not the R346T substitution, has emerged as globally dominant, likely owing to its acquisition of the Q493E substitution, also located within the RBD. This sublineage and its derivatives, particularly KP.3.1.1, now account for most SARS-CoV-2 infections ^13^.

The JN.1 cluster of sublineages occupies a unique antigenic space separate from all other prior Omicron lineages. Real-world studies of the XBB.1.5-adapted COVID-19 vaccines demonstrated robust immune response and effectiveness against XBB lineages that diminished in response to the BA.2.86 and JN.1 lineages ^14–19^. To better understand the structural and functional consequences of the antigenic shift from this new lineage cluster, and subsequent antigenic drift within it, we evaluated the biophysical and immunologic features of the prefusion-stabilized full-length SARS-CoV-2 S proteins of JN.1 and KP.2. To our knowledge, we present the first in-depth structural analysis of JN.1 and KP.2 full-length, prefusion-stabilized S immunogens, building prior reports detailing the impact of JN.1 sublineage-specific S mutations on neutralization resistance. We also assessed whether an Omicron JN.1 or KP.2-updated BNT162b2 vaccine could improve immune responses that are important for conferring protection against COVID-19 ^20^, against the JN.1 lineage and a panel of contemporary JN.1-derived sublineages, as compared to the BNT162b2 XBB.1.5 vaccine.

Both monovalent JN.1- and KP.2-adapted COVID-19 vaccines have been recommended for the 2024-2025 formula. The two lineages differ in their S amino acid sequences by 3 residues ^21^. In prior years, small genetic differences have not necessarily translated into antigenic differences. Given that two variant-adapted vaccines have been introduced in the same season, we sought to understand the impact of these few, but potentially consequential, genetic differences on biophysical, structural and immunologic attributes of vaccine-encoded JN.1 and KP.2 prefusion stabilized S proteins. The BNT162b2 JN.1 and KP.2 vaccines use the same mRNA backbone as prior approved versions of BNT162b2 and are minimally modified to contain lineage-specific sequence changes of the SARS-CoV-2 prefusion stabilized S protein that has been evaluated in previous preclinical and clinical studies. We employed a vaccine-experienced murine model, with varying schedules of prior vaccination regimens, together with a vaccine-naïve murine model, to assess humoral and cellular immunogenicity, so as to approximate the diversity of the immune-experience of the general population across all age groups ^22^. An evaluation of both structural and functional consequences of the most recent evolution of SARS-CoV-2 lineages in varied models may then elucidate the potential impact on vaccine performance.

## METHODS

### SARS-CoV-2 S(P2) and RBD Protein Expression and Purification

Protein sequences of the full-length prefusion stabilized S(P2) of the SARS-CoV-2 JN.1 lineage and KP.2 sublineage contained amino acid changes relative to the ancestral Wuhan-Hu-1 S (GenBank accession number: MN908947.3) as listed in Table S1. S(P2) contains two proline substitutions at residues 986 and 987 (K986P, V987P) and utilizes a C-terminal TwinStrep tag. The RBD constructs contain an N-terminal S protein leader peptide and coding regions from 321-527 for JN.1 and KP.2, and 327-528 for the ancestral strain, followed by a C-terminal affinity tag as indicated in Table S1. Details on S(P2) and RBD protein expression and purification can be found in the Supplementary Materials.

### Stability of Wild Type (WT), Omicron JN.1 and KP.2 FL S(P2) by Thermal Shift Assay (TSA)

Stability of FL S(P2) proteins was measured by Tycho NT.6 (NanoTemper, firmware version: 1.10.3) as described previously ^23^. The inflection temperature for each thermal melting curve reported by the Tycho NT.6 software was reported as the thermal melting temperature (T_m_) of S(P2) proteins.

### Binding Kinetics of Purified FL S(P2) Protein and RBD to Immobilized Human ACE-2-PD

FL S(P2) and RBD proteins of JN.1, KP.2 and WT strains were assessed by biolayer interferometry (BLI) binding to immobilized human ACE-2-PD, as described previously ^23^.

### Cryogenic Electron Microscopy (Cryo-EM) of JN.1 and KP.2 S(P2)

SARS-CoV-2 FL S(P2) (K986P and V987P substituted) proteins of the JN.1 and KP.2 lineages were interrogated by electronic cryogenic electron microscopy (cryo-EM) analysis following membrane extraction and purification. Grids were prepared with JN.1 and KP.2 S(P2) proteins at 5.6 and 5.8 mg/ml by applying 4 µL to Quantifoil R1.2/1.3 200 mesh gold grids that were glow-discharged for 60 seconds at 15 mA in a Pelco easiGlow cleaning system and blotted for 3 seconds with a blot force of +3 before plunging into liquid nitrogen-cooled liquid ethane using a Mark IV Vitrobot with a sample chamber maintained at 4°C and 100% humidity. Data collection was performed with a Titan Krios G2 electron microscope operated at 300 kV, controlled by EPU, and equipped with a Falcon 4i direct electron detector and Selectris energy filter configured with a 10 eV slit (ThermoFisher Scientific). A total of 7k and 9k movies were collected for JN.1 and KP.2 S(P2) protein samples with a total dose of 40.0 e^-^/Å^2^ and a −0.4 μm to −2.4 μm defocus range. Details on cryo-EM data processing and model building can be found in the Supplementary Materials and Table S3.

### Mass Spectrometry Characterization of JN.1 and KP.2 S(P2) N-linked Glycosylation

Mapping of N-linked glycosylation sites was conducted on recombinant purified JN.1 and KP.2 S(P2) as described previously ^23^. For glycosylation complexity determination, S(P2) protein digests, before and after O18 water and PNGase F treatment, were analyzed using liquid chromatography-mass spectrometry (LC-MS). The data were processed using Protein Metrics PMI-Byos Byologic PTM workflow with glycan modifications enabled. Relative levels of the various glycan forms were determined by summing the extracted ion chromatogram peak areas for all detected charge states and forms for a given isobaric glycosylation state. Relative estimation of abundance was obtained by comparing frequencies of different glycans, with an assumption that all glycopeptides have similar ionization.

### Animal Ethics

All mouse studies were performed at Pfizer, Inc. (Pearl River, NY, USA), an Association for Assessment and Accreditation of Laboratory Animal Care (AAALAC) accredited facility. Animals were observed after all procedures and injection sites were monitored following each vaccination. All procedures performed on animals were in accordance with regulations and established guidelines and were reviewed and approved by an Institutional Animal Care and Use Committee or through an ethical review process.

### Immunogenicity Studies

#### 4^th^ or 5^th^ Vaccination in BNT162b2-Experienced Mice

Monovalent XBB.1.5, JN.1, and KP.2-adapted BNT162b2 vaccines were evaluated as either a 4^th^ or 5^th^ dose in female BALB/c mice (10 per group; Jackson Laboratory). In the 5^th^ dose study, mice were vaccinated intramuscularly (i.m.) at 6-8 weeks of age with 2-doses (Day 0, 21) of original BNT162b2 WT vaccine, followed by a 3^rd^ dose booster (Day 49) of bivalent WT + Omicron BA.4/5 vaccine, a 4^th^ dose (Day 84) of the monovalent Omicron XBB.1.5 adapted vaccine, and a 5^th^ dose (Day 111) of either the monovalent XBB.1.5, JN.1, or KP.2-adapted vaccine. All vaccinations used a 0.5 µg total dose in a 50 µL volume (bivalent formulations contained equal quantities of each mRNA, 0.25 µg each). A control group of five mice received saline injections (50 µL) according to the same schedule, in place of active vaccines. Sera were collected for evaluation of pseudovirus neutralizing antibody responses prior to the 5^th^ dose (Day 111) and at the study end (Day 140). Spleens were collected from all mice at Day 140 to evaluate cell-mediated immune responses (Fig. S4A).

In the 4^th^ dose study, female BALB/c mice (10 per group; Jackson Laboratory) were first vaccinated i.m. at 6-8 weeks of age and subsequent doses were administered according to the same schedule as the 5^th^ dose study, except that the 4^th^ dose was administered on Day 64 as either the monovalent XBB.1.5, JN.1, or KP.2-adapted vaccine and there was no 5^th^ dose. As above, sera were collected for evaluation of pseudovirus neutralization responses prior to the 4^th^ dose (Day 64) and two weeks later at study end (Day 78) (Fig. S4B).

#### 2-Dose Primary Series in Naïve Mice

Female BALB/c mice (10 per group Jackson Laboratory) were vaccinated i.m. at 6-8 weeks of age on Days 0 and 21 with either monovalent XBB.1.5, JN.1, or KP.2-adapted vaccine. As described above, vaccine formulations contained a total dose level of 0.5 µg mRNA in a 50 µL volume and a control group (10 mice) received saline injections adjacent to the vaccine groups. Sera and spleens (5 mice/group) were collected 28 days after the second dose (day 49) for evaluation of pseudovirus neutralizing antibody responses and cell-mediated immune responses, respectively (Fig. S4C).

### VSV-SARS-CoV-2 S Pseudovirus Neutralization Assay

The pseudovirus neutralization assay (PNA) was defined and performed as previously described^23^. VSV-based pseudoviruses contained the S protein from the following SARS-CoV-2 lineages: WT (Wuhan-Hu-1, ancestral strain), BA.4/5, XBB.1.5, JN.1, JN.1.6,1, JN.1.7, JN.1.13.1, KW.1.1, KP.2, KP.2.2, KP.1.1, KS.1, JN.1.16.1, KZ.1.1.1, KP.2.3, KP.3, KP.3.1.1, LB.1 and XEC. Amino acid sequence substitutions relative to JN.1 for all tested pseudoviruses are provided in Fig. S5.

### Omicron XBB.1.5, JN.1 and KP.2 Antigenic Cartography

Antigenic maps were constructed using the antigenic cartography toolkit Racmacs v1.2.9 (https://acorg.github.io/Racmacs/index.html). In brief, antigenic cartography is a method to quantify and visualize neutralization data. An antigenic map takes titration data that measures strength of reactivity of a group of antisera (sera where an individual mouse has been vaccinated with a unique antigen of a particular SARS-CoV-2 lineage) against a group of antigens (different SARS-CoV-2 lineages). Antigenic mapping uses multidimensional scaling to position antigens (viruses) and sera in a map to represent their antigenic relationships. Antigenically similar strains are spatially close to one another on the map, while antigenically distinct strains are further apart. The spacing between grid lines is 1 unit of antigenic distance, corresponding to a two-fold dilution of antiserum in the neutralization assay. The maps were constructed by Racmacs package in R using 2000 optimizations, with the minimum column basis parameter set to “none.”

### T-cell Response by Flow Cytometry Assay

Murine splenocytes were stimulated *ex vivo* as previously described (*1*) with DMSO only (unstimulated) or specific amino acid (aa) peptide libraries (15aa, 11aa overlap, 1 to 2 µg/mL/peptide) representing SARS-CoV-2 S amino acid sequences. Five individual peptide pools represented the full-length S sequence of the ancestral Wuhan (WT) (JPT), BA.4/5 (JPT), XBB.1.5 (JPT), JN.1 (JPT) and KP.2 lineages (Mimotopes). Following stimulation, splenocytes were stained for CD154 (CD40L), IFN-γ, TNF-α, and IL-2 positive CD4^+^ and CD8^+^ T cells, as previously described ^23^. Samples were acquired on a 5-Laser Aurora system (Cytek ®) using SpectroFlo® software (version 3.1.2). The instrument was subject to daily quality control procedures using SpectroFlo® QC Beads per manufacturer recommendations. Acquired data files were analyzed using OMIQ® software. The T cell gating strategy is shown in Fig. S6.

### Data Availability Statement

The final full-length S(P2) cryo-EM density maps and models for the JN.1 (3-down and 1-up) and KP.2 (3-down, 1-up, and 2-up) conformations are deposited in the Electron Microscopy Data Bank (EMDB) and the Protein Data Bank (PDB) under accession codes EMD-46637 and PDB ID 9D8H, EMD-46638 and PDB ID 9D8H, EMD-46639 and PDB ID 9D8I, EMD-46640 and PDB ID 9D8J, and EMD-46641 and PDB ID 9D8K, respectively.

### Statistical Analysis

Mouse immunogenicity data were analyzed using SAS version 9.4. All statistical analyses were performed using ANOVA on log-transformed data. Comparisons were made on mouse sera across pseudoviruses at the last post-vaccination timepoint of each study with Dunnett’s test for multiple comparisons. For intergroup comparisons, the XBB.1.5 vaccine group was the reference; for intragroup comparisons (pseudoviruses within a vaccine group), the vaccine target lineage (XBB.1.5, JN.1 or KP.2) served as the reference. All tests were two-tailed. A p-value of less than 0.05 was considered statistically significant.

## RESULTS

### Enhanced ACE2 receptor binding and reduced thermostability of JN.1 and KP.2 S proteins compared to WT SARS-CoV-2 S protein

Full-length prefusion-stabilized S(P2) proteins of Wuhan-Hu-1 (WT), Omicron JN.1, and KP.2 were expressed from DNA corresponding to the adapted BNT162b2 RNA coding sequences using similar methods as previously reported^23^. All S(P2) proteins were expressed on the cell surface following in vitro transfection of Expi293F cells and showed similar binding to human ACE2 (Fig. S1A). Analysis of membrane-extracted and affinity-purified full-length S(P2) showed a single peak by size exclusion chromatography (SEC), with S predominantly cleaved into S1 and S2 subunits (Fig. 1A and Fig. S1B). Thermal shift assay (TSA) analyses showed the melting temperature (T_m_) of JN.1 (61.6 ± 0.24 ° C) and KP.2 (60.1 ± 0.19 ° C) were similar to one another (Fig. 1B) and approximately 5-7 °C lower than WT S(P2) (67.1 ± 0.17 °C) ^23^. Biolayer interferometry (BLI) analysis of purified RBDs only, which contain the receptor binding motif (RBM) critical for ACE2 binding, showed significantly increased binding of soluble JN.1 and KP.2 RBDs to fixed human ACE2 peptidase domain (ACE2-PD) (K_D_ 1.70 nM and 2.07 nM, respectively) compared to the WT RBD (K_D_ 31.3 nM) ^23^ (Fig. 1C).

**Figure 1.**
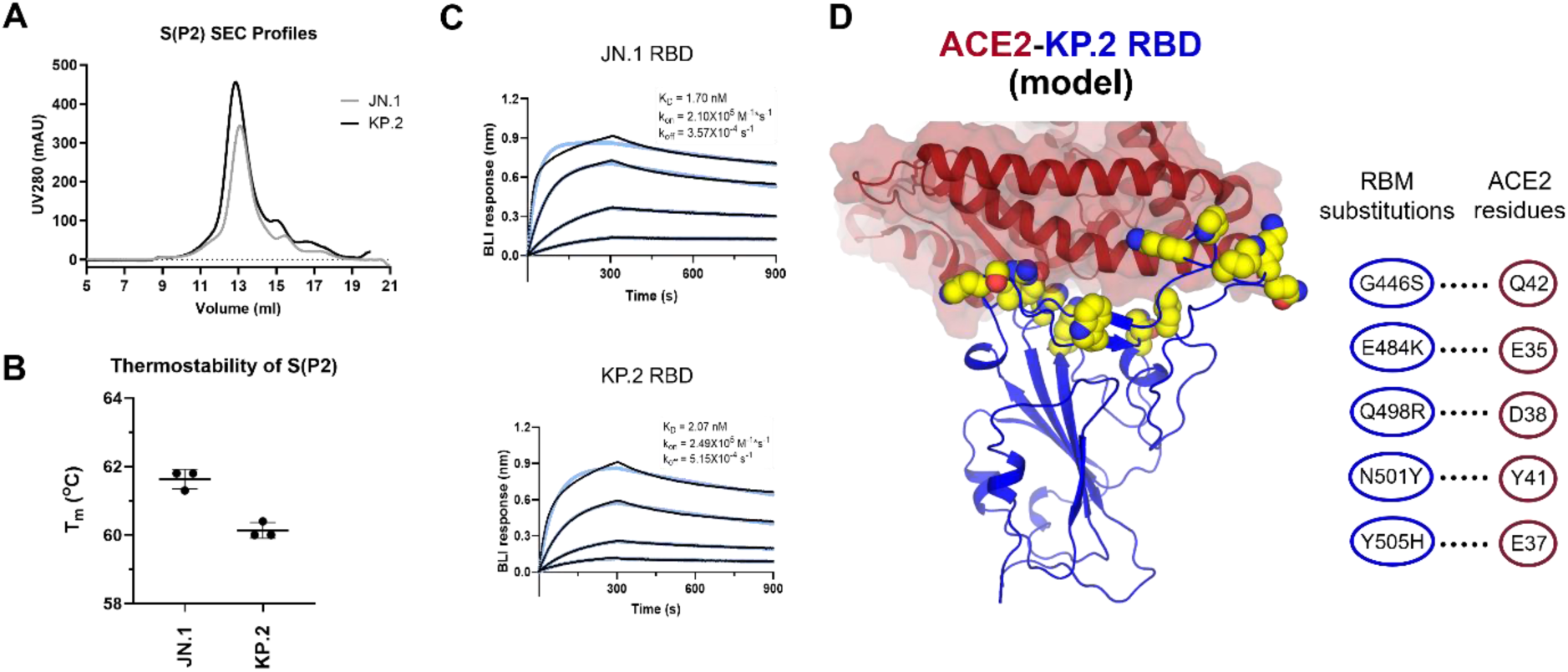
Biophysical Characteristics of SARS-CoV-2 Omicron JN.1 and KP.2 Prefusion-Stabilized Spike and Receptor Binding Domain. **a** SEC profiles of the S(P2) proteins of JN.1 and KP.2. The S(P2) proteins were purified in DDM (*n*-dodecyl-*β*-D-maltopyranoside, see Supplementary Materials). SDS-PAGE fractions from 12 mL to 15 mL of each FL S construct are shown in Fig. S1B. **b** Melting temperature (Tm) of purified S(P2) proteins determined by the inflection point of the first derivatives of protein fluorescence signals collected at 330 nm over that of 350 nm. Experiments were conducted by Tycho NT.6 and run in triplicates (n=3). **c** Biolayer interferometry (BLI) sensorgrams of RBD binding to immobilized human angiotensin converting enzyme-2 peptidase domain (ACE2-PD). The binding curves (black) were globally fit to a 1:1 Langmuir binding model (light blue). Calculated apparent K_D_, k_on_, k_off_ values are listed in the figure. **d** Crystal structure of ACE2-PD (PDB: 6M0J) is aligned to the KP.2 1-up RBD. ACE2-PD is shown in maroon ribbon with surface representation. KP.2 RBD is shown in blue cartoon with amino acid residue changes in the receptor binding motif (RBM) from WT to KP.2 shown in yellow spheres. RBM residue changes (blue circle) that enhance receptor binding via putative specific interactions (dotted lines) to ACE2 residues (maroon circle) are shown in the interaction scheme on the right.

### Acquisition of glycosylation sites within the JN.1 and KP.2 S proteins

The purified S(P2) proteins of JN.1 and KP.2 were analyzed by liquid chromatography mass spectrometry (LCMS) to identify N-linked glycosylation sites. Glycosylation mapping confirmed that JN.1 and KP.2 have conserved glycosylation sites on the S protein, despite undergoing extensive mutations compared to WT S. This is consistent with previous studies of earlier lineages, suggesting a correlation between viral fitness and glycosylation ^24^. Compared to XBB.1.5, we discovered two new glycosylation sites (Table 1) in the RBD and N-terminal domain (NTD) of JN.1 and KP.2 S, respectively. The novel RBD glycosylation site is located on N354 which is gained by the K356T substitution, whereas the novel NTD glycosylation site is a result of the H245N substitution within the peptide ALN^245^RSYLT. Analysis of the glycosylation heterogeneity confirmed that both sites are fully modified by glycans with a major composition being HexNAc(2)Hex(5) (Table S2).

**Table 1.** Emerging N-link glycosylation sites at JN.1 and KP.2 Spike proteins.

| Variant | Mutation | Peptide | % Glycosylation |
| --- | --- | --- | --- |
| JN.1 | K356T | R.FASVYAW<u>N</u>R.T | 99.8% |
|  | H245N | R.FQTLLAL<u>N</u>R.S | 99.6% |
| KP.2 | K356T | W.<u>N</u>RTRISNCVADY.S | 98.3% |
|  | H245N | R.FQTLLAL<u>N</u>R.S | 96.2% |

### JN.1 and KP.2 S(P2) cryo-EM structure

Structures of JN.1 and KP.2 full-length, prefusion stabilized SARS-CoV-2 S proteins were solved by cryo-EM to validate the conformation of the encoded antigen of both vaccines and to evaluate the structural consequences of the lineages’ hallmark mutations in the context of S(P2). Both JN.1 and KP.2 S(P2) proteins exhibited prefusion conformations, as anticipated. The cryo-EM processing employed herein sought to identify all RBD (up/down) configurations (Fig. S2) within the S trimer. Unlike the WT S(P2) protein, which was predominantly (∼80%) found in the 3-down conformation^25^, the prefusion-stabilized JN.1 S protein was observed in the 3-down (65%) conformation, and at a lower frequency in a 1-up (35%) conformations, while KP.2 S was more evenly observed in 3-down (41%) and 1-up (48%) conformations, with a small fraction in a 2-up (11%) conformation, indicating KP.2 exhibits a greater tendency than earlier variants for adopting the RBD-up conformations (Fig. 2A).

**Figure 2.**
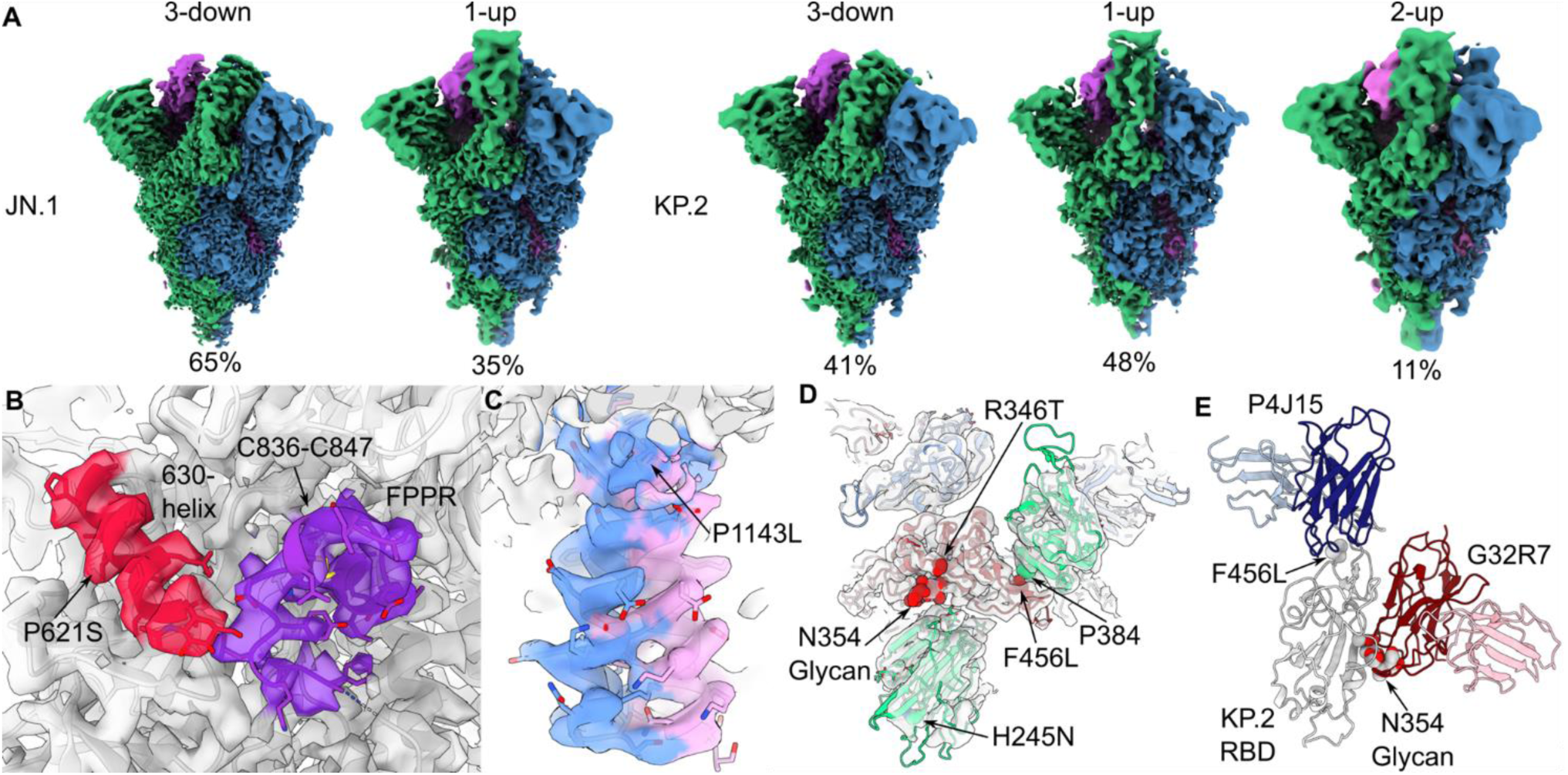
Structural and Functional Consequences of SARS-CoV-2 JN.1 and KP.2 S Protein Mutations. **a** Local resolution-filtered maps of the final 3D reconstructions are shown for each of the resolved RBD conformational states and colored by protomer. Local resolution estimations for maps shown in panels A though D are found in Figure S3. The first up RBD is colored green in both the JN.1 and KP.2 maps and the second up RBD in the KP.2 is colored blue. Proportions of particles derived from reference-based classification are expressed as a percentage below each map. **b** Magnified view of the 630-helix and fusion peptide proximal region (FPPR) from the KP.2 1-RBD-up map overlaid with the model shows the quality of the EM density in this region. **c** Magnified view of the stem from the JN.1 3-RBD-down map overlaid with the model shows the two partially occupied conformations of this helix beginning at the P1143L mutation. **d** Top view of the KP.2 1-RBD-up conformation structure shows the location of the N354 glycan due to the K356T substitution and a putative interprotomer hydrophobic interaction with P384 due to the F456L substitution. **e** Broad monoclonal antibodies (mAbs) G32R7 (PDB: 7N65) and P4J15 (PDB: 8PQ2), which bind early omicron lineages, are aligned to the KP.2 1-RBD-up. The proximity of these epitopes and the K356T and F456L substitution-induced surface alterations implicate these changes with evasion of early immunity.

In both JN.1 and KP.2 S, the C3-symmetric, 3-down conformation of the RBD core is resolved at ∼4Å, whereas the receptor binding motif (RBM), comprising residues 450-490, were absent at high resolution. In the 1-up conformations, the RBM was best resolved for the down-RBD adjacent to the up-RBD (Fig. S3). The RBM of the down-RBD, adjacent to the up-RBD, was poorly resolved, though it contained high-resolution features for the core. In contrast, very little of the second up-RBD was seen in the 2-RBD-up KP.2 structure (Fig. S3).

Structural changes for both lineages were likely caused by two amino acid substitutions shared by JN.1 and KP.2—P621S and P1143L—that differentiate them from XBB.1.5. The P621 residue is in a region of the S subdomain 2 (SD2) called the 630 loop that is unstructured in full-length, 2P-stabilized XBB.1.5 and Wuhan S protein cryo-EM densities. Despite being situated in this unstructured region, the P621S substitution leads to a well-resolved alpha helix (Fig. 2B). The structured region is found in every protomer of the five solved structures of the JN.1 and KP.2 S proteins. In the context of the 1-up RBD structures of JN.1 and KP.2, a region adjacent to the 630 helix—residues 829-848 or the fusion peptide proximal region (FPPR) of the adjacent protomer—is also well-resolved (Fig. 2B, Fig. S3). Though lower in resolution, there is a clear density, including a disulfide bond between C836 and C847. As a result, the structured FPPR interacts with the 630-helix from the adjacent protomer, contributing to an enhanced interprotomer stability which may reduce S1/S2 shedding in the context of the RBD-up conformation. The P1143 residue is located near the N-terminus of the stem helix. The cryo-EM density revealed two discrete conformations of the stem for both JN.1 and KP.2 S. The alternative poses were best resolved in the JN.1 3-down, C3-symmetry reconstruction (Fig. 2C, Fig. S3) and diverge near the mutated proline residue.

Two additional mutations, H245N and K356T, differentiate JN.1 and KP.2 from XBB.1.5 and earlier lineages, and are shared with the parental BA.2.86 lineage. These mutations have yielded novel glycosylation sites that were fully glycosylated in the resolved structures (Fig. 2D, Tables 2 and S1). However, H245N within the NTD is located in a low-resolution region of the EM densities of the two JN.1 and three KP.2 structures. As a result, the possible glycosylation modification of H245N is absent from the cryo-EM map reconstruction. The K356T mutation led to glycosylation at N354 in JN.1, KP.2 and previous variants harboring the mutation ^26,27^. This glycan is located near the interface of the RBD and the NTD of an adjacent protomer (Fig. 2D).

Overall, JN.1 and KP.2 S proteins differ by three residues: R346T and F456L in the RBD and V1104L in the S2 domain. R346T has been observed in several earlier lineages and may impact antibodies recognizing the RBD class 3 epitope ^28^. The V1104L mutant residue exhibited density consistent with the additional methylene group and no discernable structural changes. F456L is located in the RBM and in a relatively low-resolution region of the map. Of note, the RBD is best resolved in the down conformation adjacent to the up conformation and the mutant L456 is located at the protomer interface (Fig. 2D, Fig. S3).

### BNT162b2 JN.1- and KP.2-adapted vaccines neutralizing responses to JN.1 sublineages in a vaccine-experienced mouse model

BNT162b2 JN.1- and KP.2-adapted vaccines were evaluated in two murine studies that examined booster immunogenicity in a BNT162b2-experienced immune setting. To approximate the immune background of a vaccinated human population, a vaccine-experienced animal model was generated by vaccinating mice with all licensed BNT162b2 vaccines sequentially spanning 2021 to the present [Original (2-doses), Bivalent Original+BA.4/5 (1-dose), XBB.1.5 (1-dose)]. BNT162b2 XBB.1.5, JN.1, or KP.2-adapted vaccines were administered as a 5^th^ dose to female BALB/c mice 27 days following the XBB.1.5 vaccination (4^th^ dose) (Fig. S4A). Sera were collected prior to and one month following administration of the 5^th^ dose for assessment of pseudovirus neutralization against contemporary JN.1 lineages (JN.1, JN.1.16.1, KP.2, KP.2.3, KP.3, KP.3.1.1, LB.1, and XEC).

As a 5^th^ dose, JN.1- and KP.2-adapted vaccines elicited much higher 50% geometric mean neutralizing titers (GMTs) against JN.1 and all JN.1 sublineages tested, as compared to the XBB.1.5 vaccine (Fig. 3A). The JN.1 and KP.2 vaccine responses were 3-to-4 times and 7- to-10 times higher, respectively, compared to the XBB.1.5 vaccine group, including against the globally prevalent KP.3.1.1 sublineage and the rapidly rising XEC sublineage (Fig. 3B). In XBB.1.5 vaccinated animals, neutralizing responses against the antigenically distant JN.1 lineage were approximately 20-fold lower than against the vaccine-matched lineage, XBB.1.5 (Fig. 3A). In contrast, sera from JN.1- and KP.2-vaccinated mice neutralized all JN.1 sublineages with similar potency, indicating a broadly robust and cross-protective immune response. From pre- to post-5^th^ dose, the JN.1-adapted vaccine boosted neutralizing responses (GMT fold rise (GMFR)) against JN.1 and KP.2 by 3.5- and 3.9-fold, respectively. The KP.2-adapted vaccine increased neutralizing responses by 5.9-fold and 7.4-fold against the JN.1 and KP.2 lineages, respectively, as compared to the XBB.1.5 vaccine (Fig. S7). The XBB.1.5-adapted vaccine elicited the lowest GMFRs against JN.1 and KP.2 (0.9 and 1.8, respectively).

**Figure 3.**
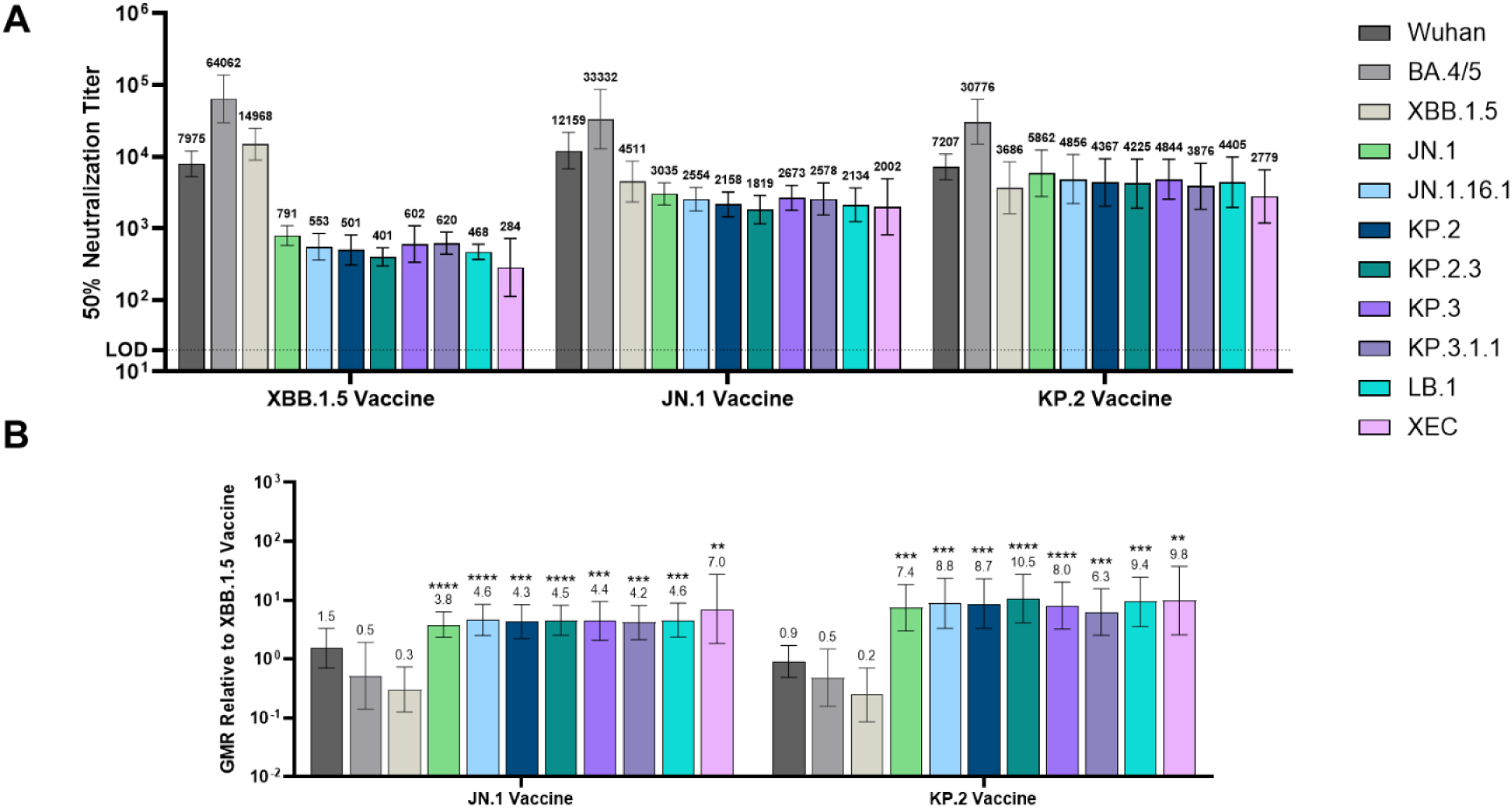
Pseudovirus Neutralization Elicited by BNT162b2 XBB.1.5, JN.1 And KP.2-Adapted Vaccines Administered to Vaccine-Experienced Mice. Female mice were immunized i.m. according to Fig. S4A on Days 0 and 21 with the BNT162b2 Wuhan (WT), on Day 49 with the bivalent BNT162b2 (WT + BA.4/5), on Day 84 with BNT162b2 XBB.1.5, and on Day 111 with the BNT162b2 XBB.1.5, JN.1, or KP.2 vaccine. One month post-5^th^ dose, sera were collected from the terminal bleed and neutralizing antibody responses were measured against a panel of 11 pseudoviruses that included the Wuhan (WT) reference strain and Omicron lineages BA.4/5, XBB.1.5, JN.1, JN.16.1, KP.2, KP.2.3, KP.3, KP.3.1.1, LB.1, and XEC. **a** The number above each bar indicates the 50% neutralizing geometric mean titer (GMT) with 95% CI of 10 mice per vaccine group. **b** The geometric mean ratio (GMR) is shown as the ratio of the BNT162b2 KP.2 or JN.1 vaccine GMT to the BNT162b2 XBB.1.5 vaccine GMT of the corresponding pseudovirus. The number above each bar indicates the GMR with 95% CI. The limit of detection (LOD) is the lowest serum dilution, 1:20. Asterisks indicate statistical significance of pseudovirus GMR relative to the corresponding pseudovirus in the monovalent XBB.1.5 vaccine group. ****p<0.0001, ***p<0.001, ** p<0.01, * p<0.05.

In a separate study, BNT162b2 XBB.1.5, JN.1, or KP.2 vaccines were administered as a 4^th^ dose to female BALB/c mice two weeks after the 3^rd^ dose (BNT162b2 bivalent WT + BA.4/5) (Fig. S4B). As observed in the 5^th^ dose study, sera from JN.1- and KP.2-vaccinated mice neutralized a broad panel of JN.1 sublineages with similar potency (Fig. S8A). Here again, the JN.1- and KP.2-adapted vaccines elicited significantly (p <0.05) higher neutralizing responses against JN.1 sublineages, on the order of 2-to-4-fold and 3-to-7-fold, respectively, than the XBB.1.5 vaccine. GMFRs from pre-to-post 4^th^ dose were also similar between the JN.1 and KP.2 vaccines, as compared to the XBB.1.5-adapted vaccine and similar to the magnitude of rise observed when the vaccines were given as a 5^th^ dose (Fig. S8B). Despite the variations in prior vaccination regimens, the JN.1 and KP.2-adapted vaccines elicited consistent neutralization trends and significantly improved immunogenicity against contemporary JN.1 lineage pseudoviruses, including against the more recently dominant sublineages (*i.e.*, KP.3.1.1 and XEC), as compared to the XBB.1.5 vaccine.

### BNT162b2 JN.1 and KP.2-adapted vaccines neutralizing responses to JN.1 sublineages in naïve mice

The Omicron XBB.1.5, JN.1 and KP.2 vaccines were administered on Days 0 and 21 to naïve female BALB/c mice as a primary series (Fig. S4C). Sera were collected one month after the 2^nd^ dose and tested against the pseudovirus panel used in the vaccine experienced studies described above. Although the BNT162b2 XBB.1.5 vaccine induced robust neutralizing responses against the XBB.1.5 sublineage, it failed to elicit similar neutralizing titers against the antigenically distant JN.1 lineage and sublineages (Fig. 4A). Overall, the JN.1 and KP.2-adapted vaccines elicited significantly higher neutralizing responses against JN.1 and other relevant JN.1 sublineages, compared to the XBB.1.5-adapted vaccine (Fig. 4), by an order of magnitude greater than the differences observed in the vaccine-experienced models. Neutralizing GMTs in the JN.1 and KP.2 vaccine groups were 9-to-14-fold and 15-to-29-fold higher against JN.1 and JN.1 sublineages, respectively, as compared to the XBB.1.5 vaccine group (Fig. 4B). Although KP.2-adapted vaccine-elicited GMTs trended 2-to-3-fold higher than JN.1 vaccine-elicited GMTs, these differences were not statistically significant (Fig. 4B).

**Figure 4.**
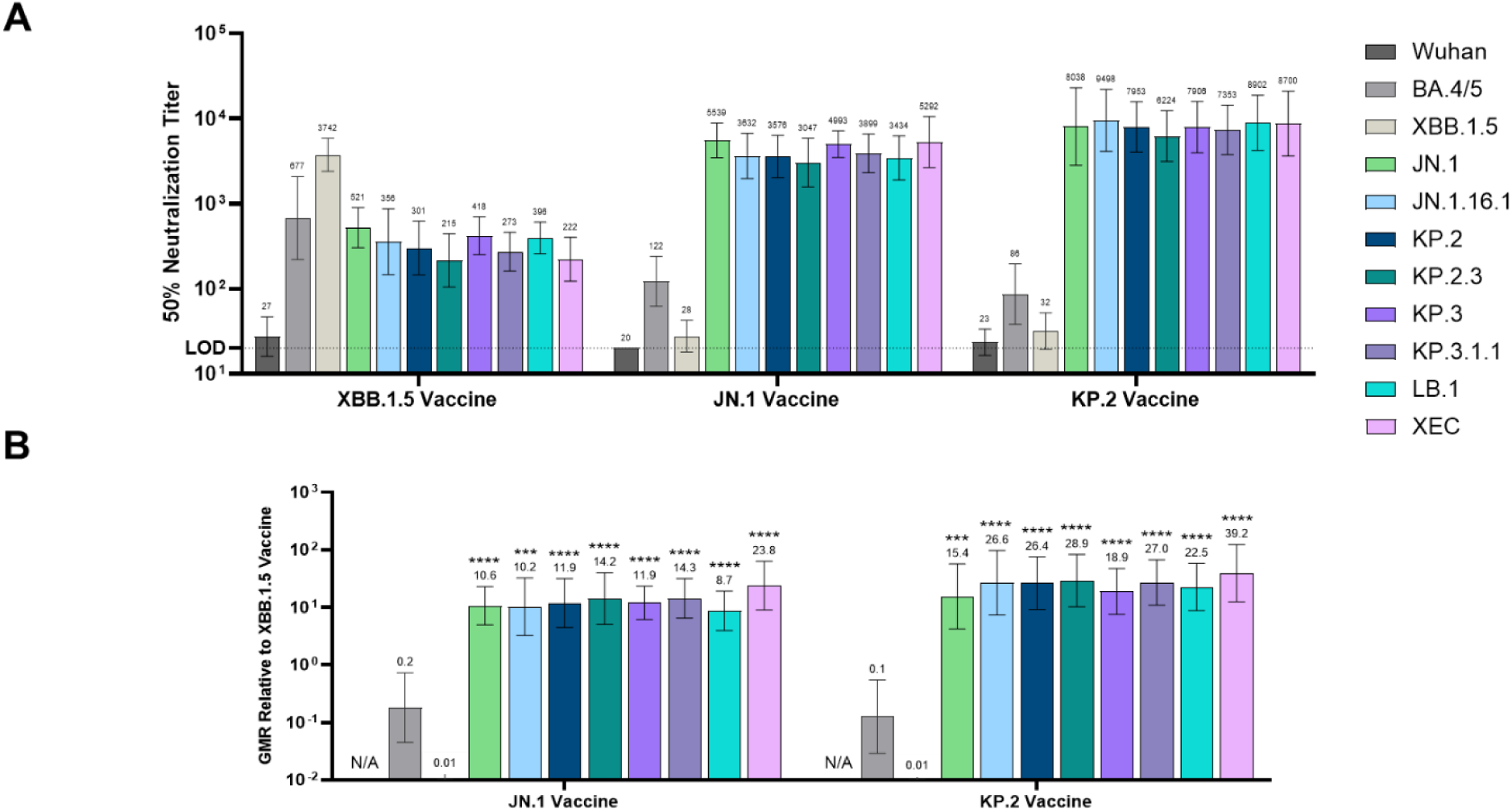
Pseudovirus Neutralization Elicited by BNT162b2 XBB.1.5, JN.1 and KP.2-Adapted Vaccines Administered as a Primary Series to Naïve Mice. Female mice were immunized i.m. according to Fig. S4A on Days 0 and 21 with the BNT162b2 XBB.1.5, JN.1, or KP.2-adapted vaccine. Sera were collected from the terminal bleed (Day 49) and neutralizing antibody responses were measured against a panel of 11 pseudoviruses that included the Wuhan (WT) reference strain and Omicron lineages BA.4/5, XBB.1.5, JN.1, JN.16.1, KP.2, KP.2.3, KP.3, KP.3.1.1, LB.1, and XEC. **a** The number above each bar indicates the 50% neutralizing geometric mean titer (GMT) with 95% CI of 10 mice per vaccine group. **b** The geometric mean ratio (GMR) is shown as the ratio of the BNT162b2 KP.2 or JN.1 vaccine GMT to the BNT162b2 XBB.1.5 vaccine GMT of the analogous pseudovirus. The number above each bar indicates the GMR with 95% CI. The limit of detection (LOD) is the lowest serum dilution, 1:20. Asterisks indicate statistical significance of pseudovirus GMR relative to the analogous pseudovirus in the monovalent XBB.1.5 vaccine group. ****p<0.0001, ***p<0.001, ** p<0.01, * p<0.05. N/A = Not available as both values to calculate GMR are at limit of detection.

### Mapping of neutralizing antibody responses reveals antigenic shifts and drifts of SARS-CoV-2 Omicron lineages

To investigate the relative antigenic differences among lineages, as reflected in variant-adapted vaccine humoral immunogenicity, serum neutralizing titers from XBB.1.5-, JN.1- and KP.2-adapted BNT162b2 vaccinated mice (vaccine-experienced and naïve) described above, were used to generate antigenic maps of contemporary SARS-CoV-2 lineages, relative to one another and to the ancestral Wuhan-Hu-1 (WT) strain (Fig. 5). In the map of vaccine-elicited sera from naïve mice, JN.1 and all JN.1-derived sublineages lie within 2 antigenic units (1 unit equals 2-fold change in neutralization titer) from each other, suggesting a high antigenic similarity and indicative of limited antigenic drift within the JN.1 cluster thus far (Fig. 5A). In contrast, JN.1 and JN.1-derived sublineages were found to lie more than 4 antigenic units away from the original WT strain, and previously dominant lineages BA.4/5 and Omicron XBB.1.5, indicating a major antigenic shift from earlier SARS-CoV-2 lineages. Sera from vaccine-experienced mice yielded similar spatial relationships (Fig. 5B); however, these data are likely confounded by the influence of cross-reactive antibodies recognizing conserved epitopes. Antigenic maps generated in a naïve background more accurately reflect the true antigenic differences between virus strains or species. The cartographies of both backgrounds, however, clearly demonstrate that evolution toward JN.1 lineages marks a major antigenic shift from prior dominant lineages, such as those belonging to the XBB cluster. The distance of JN.1 lineages from XBB.1.5 is even greater than the latter is from its epidemiologically dominant predecessor, BA.4/5. However, the JN.1 lineages, particularly the parental JN.1 and its derivative KP.2, occupy proximal or overlapping coordinates of the antigenic maps, which is consistent with the similarity in neutralizing activity elicited by both vaccines against the entire panel of JN.1 sublineages.

**Figure 5.**
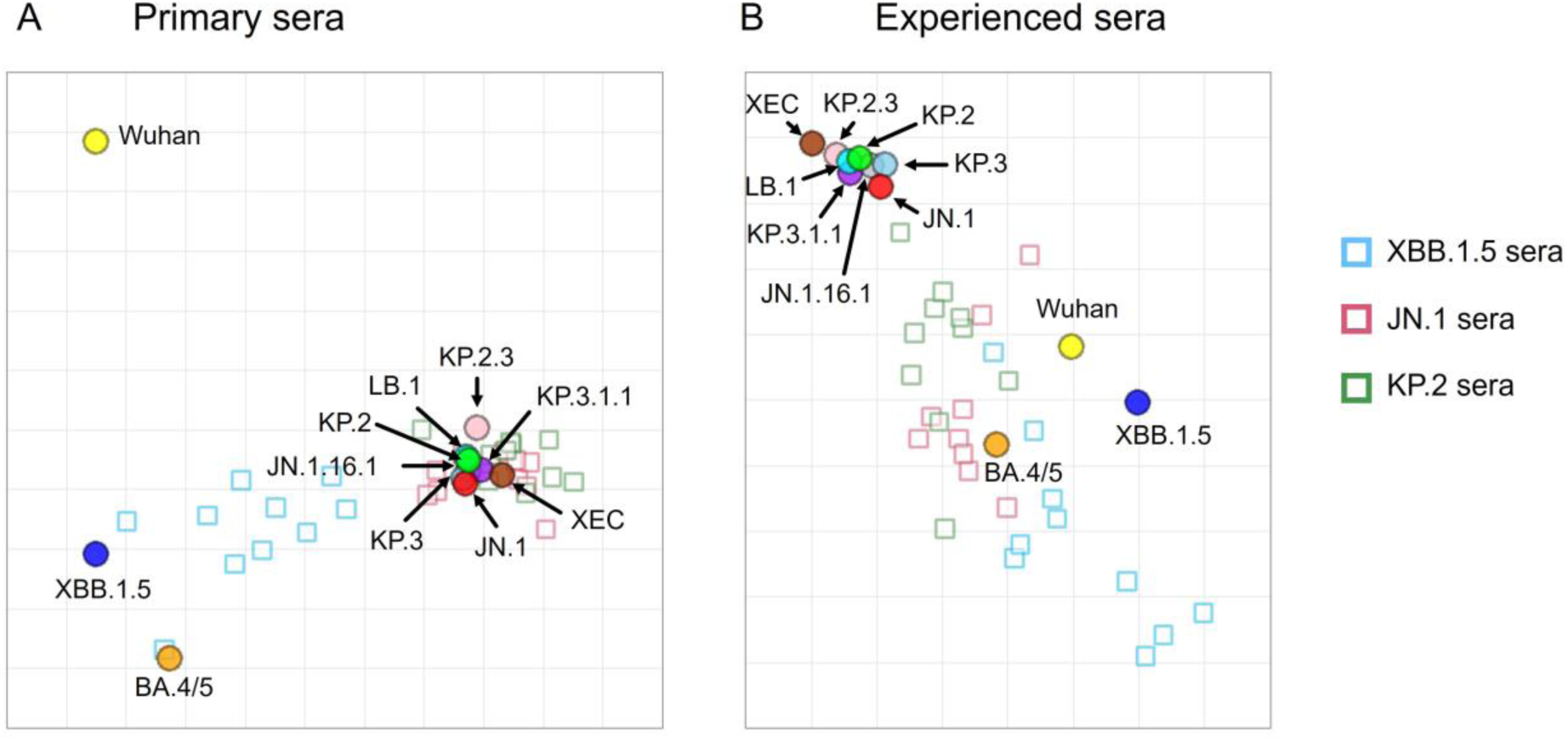
Antigenic Map of SARS-CoV-2 Omicron JN.1 Lineages Relative to Lineages Contained in Prior BNT162b2 Lineage-Adapted Vaccines. **a-b** Antigenic map visualizes cross-reactivity among a panel of 11 SARS-CoV-2 lineages showing **(a)** three groups of post-vaccination sera from naïve mice and **(b)** three groups of post-vaccination sera from vaccine-experienced mice. SARS-CoV-2 lineages are shown as circles and sera are indicated as squares. Each square corresponds to sera of one individual mouse and is colored by the vaccine that mouse received (BNT162b2 XBB.1.5, JN.1, or KP.2). Antigenic distance is represented in both horizontal and vertical axes. Each square in the matrix represents 1 antigenic unit, which reflects a two-fold difference in neutralization titer. The points that are more closely together reflect higher cross-neutralization and are therefore antigenically more similar.

### BNT162b2 JN.1- and KP.2-adapted vaccines induce comparable S-specific CD4^+^ and CD8^+^ T cell responses in BNT162b2-experienced and naïve mice

In both vaccine-experienced and naïve mice, spleens collected one-month following final vaccination were interrogated for S-specific T cell cytokine responses using a flow cytometry-based intracellular cytokine staining (ICS) assay (Fig. S6). Peptide pools representing the full-length S protein from the Wuhan (WT) virus, BA.4/5, XBB.1.5, JN.1 and KP.2 variants were used to assess lineage-specific CD4^+^ and CD8^+^ T cell responses *ex vivo*.

In vaccine-experienced mice, the XBB.1.5, JN.1 and KP.2 vaccines induced overall high frequencies of S-specific CD4^+^ and CD8^+^ T cells as compared to the saline control (Fig. 6, Fig. S9). Mice administered the 5^th^ dose (Fig. S4A) of KP.2 or JN.1-adapted vaccine elicited similar frequencies of IFN-γ^+^, TNF-α^+^, and IL-2^+^ CD4^+^ T cells, as compared to the XBB.1.5 vaccine (Fig. 6A-C). Within each vaccine group the frequencies of cytokine expressing CD4^+^ T cells were similar across all the lineages tested, indicating that the T cell responses were highly cross-reactive. Similarly, for the cytokine expressing (IFN-γ^+^ or TNF-α^+^ or IL-2^+^) CD8^+^ T cells, the KP.2 and JN.1-adapted vaccine elicited comparable frequencies as the XBB.1.5-adapted vaccine (Fig. 6D-F). Within each vaccine group, individual lineage-specific T cell frequencies varied slightly; however, these variations did not impact the overall magnitude of the T cell response, and each vaccine was capable of inducing robust T cell cytokine responses against all the lineages analyzed. The same was true for polyfunctional (IFN-γ^+^ TNF-α^+^ IL-2^+^) CD4^+^ (Fig. S9A) and CD8^+^ T cells (Fig. S9B).

**Figure 6.**
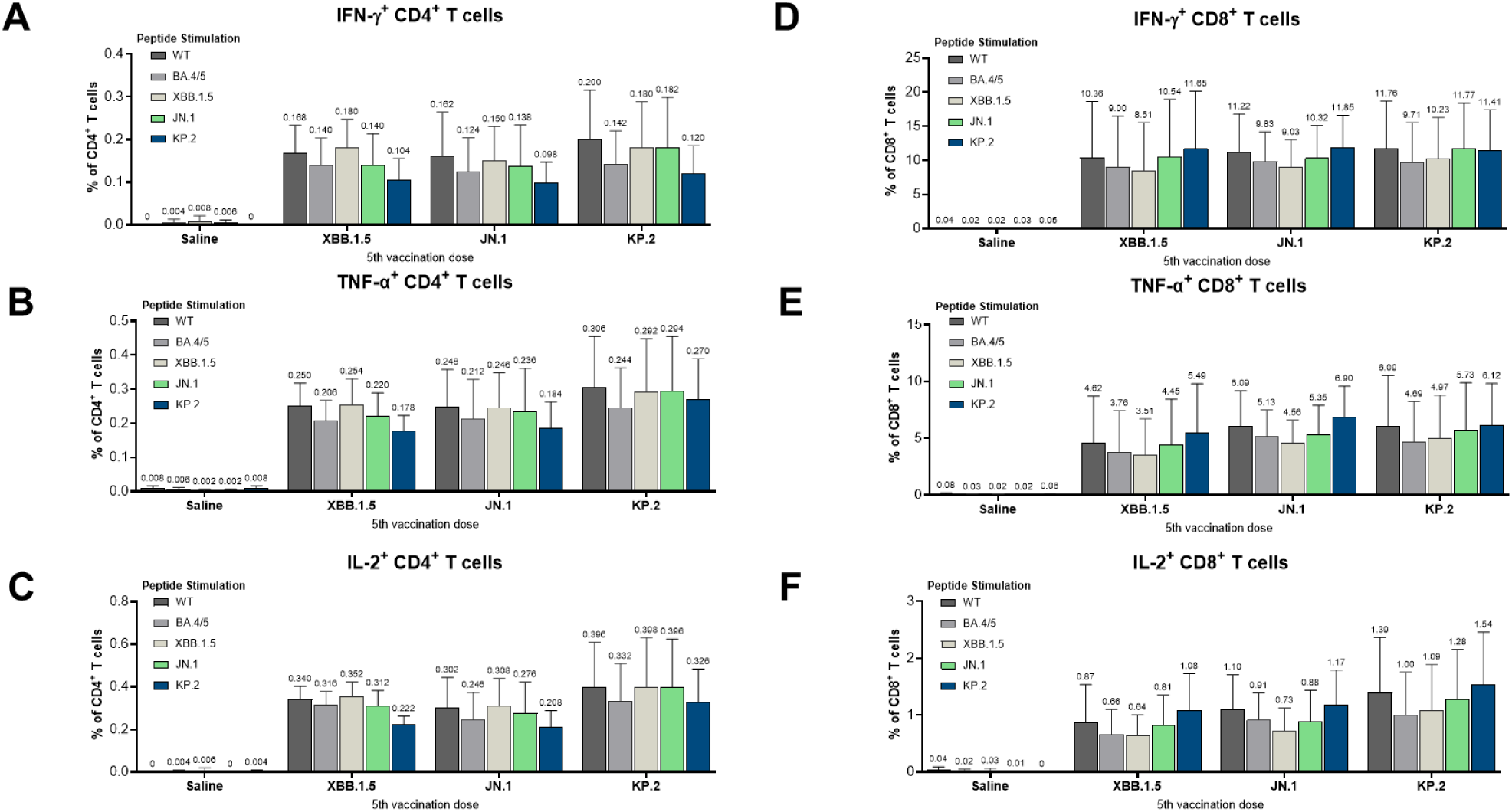
T Cell Mediated Immune Response Elicited by BNT162b2 XBB.1.5, JN.1 and KP.2-Adapted Vaccines Administered to Vaccine-Experienced Mice. One-month after the 5^th^ dose of BNT162b2 variant-adapted vaccine (XBB.1.5, JN.1, or KP.2), S-specific splenocytes (n=5/group) were characterized by a flow cytometry-based intracellular cytokine staining (ICS) assay. All samples were stimulated *ex vivo* with S peptide pools from the WT reference strain and Omicron BA.4/5, XBB.1.5, JN.1, and KP.2 sublineages. **a-f** Graphs show the frequency of CD4^+^ T cells expressing (**a)** IFN-γ, **(b)** TNF-α, and **(c)** IL-2, and the frequency of CD8^+^ T cells expressing **(d)** IFN-γ, **(e)** TNF-α, **(f)** IL-2 in response to stimulation with each peptide pool across vaccine groups. Bars depict mean frequency + SEM.

In naïve mice, the trends for both CD4^+^ (Fig. 7A-C, Fig. S10A) and CD8^+^ (Fig. 7D-F, Fig. S10B) T cell responses elicited by the JN.1 and KP.2 vaccines, with respect to cross-reactivity and similarity to one another and to the XBB.1.5 vaccine, were generally consistent with those observed in the vaccine-experienced study. In both models, minimal IL-4 secreting CD4^+^ T cells were detected (data not shown), particularly in relation to those secreting IFN-γ, TNF-α and IL-2, indicating a strong bias toward a Th1 over a Th2 cytokine profile, consistent with character of the T cell responses elicited by the original and earlier versions of BNT162b2 ^23,25^.

**Figure 7.**
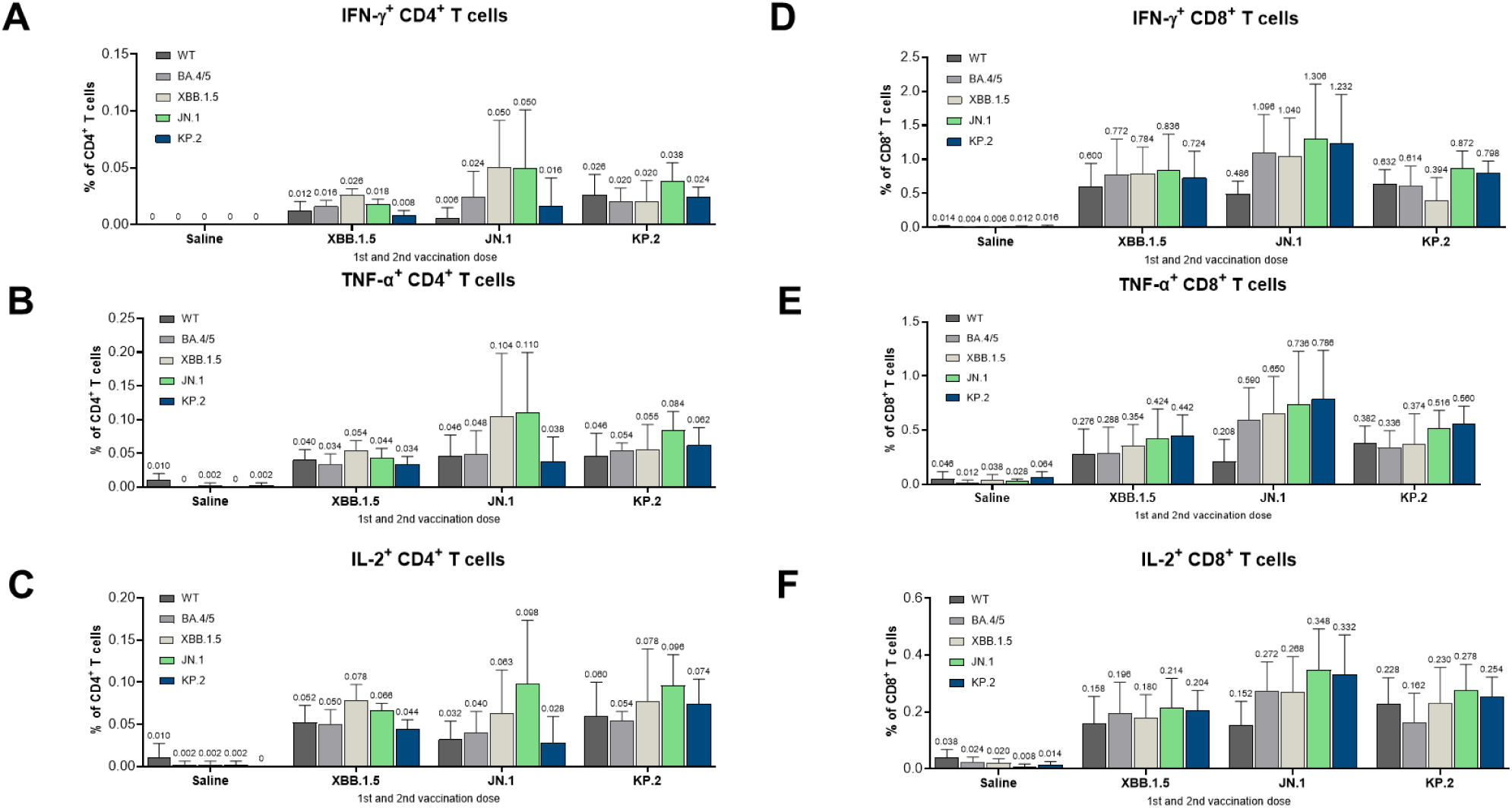
T Cell Mediated Immune Response Elicited by BNT162b2 XBB.1.5, JN.1 and KP.2-Adapted Vaccines Administered to Naïve Mice. At one-month post-second dose of BNT162b2 variant-adapted vaccine (XBB.1.5, JN.1, or KP.2) (completion of primary series), S-specific splenocytes (n=5/group) were measured by intracellular cytokine staining (ICS) assay. All samples were stimulated *ex vivo* with S peptide pools from the WT reference strain and Omicron BA.4/5, XBB.1.5, JN.1, and KP.2 sublineages. **a-f** Graphs show the frequency of CD4^+^ T cells expressing **(a)** IFN-γ, **(b)** TNF-α, and **(c)** IL-2, and the frequency of CD8^+^ T cells expressing **(d)** IFN-γ, **(e)** TNF-α, **(f)** IL-2 in response to stimulation with each peptide pool across vaccine groups. Bars depict mean frequency + SEM.

## DISCUSSION

The dynamic epidemiologic landscape of SARS-CoV-2 presents a unique challenge for the development of periodically updated COVID-19 vaccines. Since its emergence, SARS-CoV-2 has evolved quickly and broadly, comparable to other respiratory viruses that account for a large burden of disease, such as influenza ^29^. The periodicity of disease incidence, though, has not aligned with the more predictable seasonality of other respiratory viral infections ^30,31^. These attributes pose challenges for vaccine manufacturers and regulatory authorities alike in making decisions about what lineages to target in vaccine updates and when to introduce those updated vaccines into the general population, to make the greatest impact in reducing the burden of COVID-19 morbidity and mortality.

The FDA recently approved the 2024-2025 formula of BNT162b2 for individuals 12 years of age and older and granted emergency use authorization (EUA) for individuals 6 months through 11 years of age in the US ^32^, following a recommendation to mRNA vaccine manufacturers to target the KP.2 lineage in the vaccine update ^11^. The genetic drift of lineages, and questions around the relative antigenic proximity to future lineages that may emerge in the Northern Hemisphere Winter of 2024/2025, has prompted slightly different recommendations from regulatory authorities on the composition of a vaccine update ^9–11^. As such, both JN.1- and KP.2-adapted formulas of the BNT162b2 vaccine have been approved and authorized in different countries. The nonclinical evaluation of both formulas in multiple immune backgrounds against a broad panel of immunologically and epidemiologically relevant contemporary lineages, summarized here, has enabled an informed decision-making process in the development of these variant-adapted vaccines.

JN.1 differs from most of its descendant lineages by 1 to 5 amino acid residues in the S protein, which is in the same range of difference that separated XBB.1.5 from other dominant XBB sublineages ^33^. As the XBB.1.5 vaccine conferred cross-protective immunity against other sublineages within the XBB lineage cluster ^15–19,34^, we hypothesized that JN.1 and KP.2-adapted vaccines would be similarly effective in generating broad immune responses against contemporary JN.1 sublineages. Virus neutralizing antibody responses trend closely with protection from COVID-19 disease ^20,35,36^, and therefore have been a useful surrogate for COVID-19 vaccine performance against emerging lineages. The degree to which those lineages escape vaccine elicited neutralization has generally served as a metric of how well updated vaccines will protect against COVID-19.

To elucidate potential molecular mechanisms of immune escape, we sought to first characterize the JN.1 and KP.2 full-length prefusion stabilized S(P2) proteins and RBDs. Our data revealed the unique molecular features of JN.1 and KP.2 S(P2) gained from the extensive mutations during viral evolution. The S(P2) proteins of both JN.1 and KP.2 have lower T_m_ as compared to the WT and XBB.1.5 S(P2) proteins ^23^. This finding is consistent with the idea that SARS-CoV-2 may be evolving toward gradually reduced thermostability, which, along with the increased population of S in RBD-up conformations, could contribute to increased infectivity and transmissibility, and account for the epidemiologic growth advantage of these lineages relative to their predecessors. When assessing ACE2 binding affinity of the new JN.1 lineages, the RBDs of JN.1 and KP.2 have inherited several mutations also contained in XBB.1.5 that enhance receptor binding, such as G446S, Q498R, N501Y and Y505H in the RBM. The additional E484K substitution acquired by JN.1 and KP.2 may contribute to further enhancement of receptor binding via a salt bridge with E35 from ACE2 (Fig. 1D). This likelihood is supported by data that shows JN.1 and KP.2 RBD bind to ACE2 with a higher potency, as compared to the WT RBD^23^.

Few cryo-EM structures of SARS-CoV-2 S proteins have been resolved with more than one RBD-up conformation and not bound to ACE2 or antibodies ^37^. KP.2 exhibits a greater propensity to occupy the RBD-up conformation than does JN.1, despite these lineages differing at only three residues in the S protein. We reason that inter-RBD interactions conferred by F456L substitution may stabilize the RBD up conformation. In the 1- and 2-up KP.2 structures, L456 of the down-RBD is in the immediate vicinity of P384 of the up-RBD, and likely participates in hydrophobic interactions (Fig. 2D, Fig. S3). The FPPR is only well-resolved in the protomer bearing the up conformation in the 1-up structures. The improved resolution of this feature in the KP.2 versus the JN.1 structure may be due in part to the stabilization imposed by the inter-RBD contact mediated by L456, especially as RBD dynamics have been implicated in interactions within this distal region ^38,39^.

Previous non-prefusion-stabilized structures of the Omicron BA.1 lineage revealed that the 630-loop was well resolved in the down conformation ^40^, like the D614G and other earlier lineages ^38,39^. The P612S mutation may play a role similar to D614G by stabilizing the FPPR via a structured 630-loop, thereby enhancing the stability of the cleaved trimer and bolstering interactions between protomers, as well as between S1 and S2; however, doing so in the setting of the up conformation, like the Gamma variant ^38^.

The P1143 residue is located at the N-terminus of the stem helix. The secondary amide between the side chain and the main chain renders this residue unable to donate an upstream hydrogen bond. Conversely, L1143, a new mutation acquired by both JN.1 and KP.2 can capitalize on this hydrogen bonding interaction, where this additional secondary structure is likely to enforce an alternative trajectory of the stem helix. The interior facing residues of the stem are the target of weak, broadly neutralizing antibodies and the P1443L mutation may, therefore, represent an initial step toward stabilizing a conformation of the stem helix that occludes this epitope, yet is not detrimental to attaining the postfusion conformation.

Neutralizing antibodies that are overrepresented at the population level are key drivers of antigenic drift, as most that have conserved gene usage and stereotyped epitope recognition have been rendered ineffective by point mutations ^41^. For example, P4J15 is a rare antibody that recognizes the class 1 epitope and has maintained breadth from the original Wuhan strain through XBB.1 ^42^. However, the F456L mutation, present in KP.2 and located within the P4J15 epitope, likely impacts its binding (Fig. 2E). The N364 glycan is similarly located in a region recognized by broadly neutralizing antibodies, such as G32R7 (Fig. 2E). This region is known as the RBD class 3 or the RBD1 competition cluster, and represents the largest memory B cell epitope cluster among convalescent patients infected at the start of the pandemic ^28^. The mutations found in KP.2 likely perturb the epitopes of the few remaining neutralizing antibodies elicited by the Wuhan strain and underpin much of its fitness and immune escape among the SARS-CoV-2-experienced population.

As nonclinical data have closely aligned with clinical responses in prior cycles of variant-adapted vaccine updates ^14^, we assessed the immunogenicity of BNT162b2 JN.1 and KP.2 vaccines against JN.1 and contemporary circulating sublineages of epidemiologic relevance in BNT162b2-experienced and naïve mice. As most adults have developed hybrid immunity from a mixture of prior infections and vaccinations, recapitulating the collective history of antigenic exposures has become an increasingly complex endeavor, particularly with the successive emergence of new lineage clusters each year. Nonetheless, models that account for prior exposure history provide a more relevant model for evaluating vaccine responses in immune experienced populations. To understand the impact of varying immune backgrounds on variant-adapted vaccine immunogenicity, we assessed vaccine-elicited immune responses in studies with differing compositions and schedules of prior vaccinations. The most parsimonious model that maintains immunologic relevance is likely to streamline preparatory activities for future vaccine updates.

Prior preclinical data that supported FDA approval of the BNT162b2 XBB.1.5 vaccine in 2023 showed that vaccine-elicited neutralizing responses against the vaccine-matched XBB.1.5 and related lineages (*e.g*. XBB.1.16, XBB.2.3) were 4-to-5 times higher than responses elicited against the same lineage by the previously authorized vaccine (bivalent WT + Omicron BA.4/5) ^23^. Similarly, in the present studies, JN.1 and KP.2-adapted vaccines, elicited neutralizing titers against JN.1 sublineages that were consistently improved over BNT162b2 XBB.1.5 vaccine responses. Overall, there was a 3-to-10-fold improved response over the XBB.1.5 vaccine in BNT162b2-experienced mice across two studies, depending on the vaccine formulation and the sublineage tested. JN.1 and KP.2 vaccines both elicited responses that effectively neutralized all 16 JN.1 pseudoviruses evaluated, including those sublineages that acquired advantageous R346T and F456L substitutions (*e.g.*, JN.1.16.1, KP.2), and a next generation of sublineages that have an additional advantageous deletion at the S31 position (*e.g.*, KP.2.3, LB.1) ^43^. KP.3 and its most dominant KP.3.1.1 sublineage, as well as the rapidly emerging recombinant XEC lineage ^13^, were also evaluated and neutralized by JN.1 and KP.2 vaccines in the studies reported here.

As multiple animal models are employed in the evaluation of COVID-19 vaccines, many still frequently perform primary series studies in naïve animals ^44–48^, which does not account for the current immune experience of most adults. Even when vaccine-experienced animal studies are employed to better recapitulate the antigenic exposures of the general population, there remains a lack of consensus on how those studies should be designed. To inform investigations into the influence of varied immune backgrounds, we assessed JN.1 and KP.2 vaccine immunogenicity against two different vaccine-experience regimens, varying the prior vaccine composition and schedule. Across both studies, we observed a similar trend in neutralizing antibody responses for both JN.1 and KP.2 vaccines, regardless of prior BNT162b2 XBB.1.5 vaccine exposure or timing between vaccine administrations. Mice boosted with JN.1 or KP.2-adapted vaccines as a 4^th^ dose elicited greater immunity against JN.1 and a broader panel of JN.1 sublineages, consistent with findings of the 5^th^ dose study. These findings indicate that the exact sequence and scheme of prior antigenic exposures may not affect trends in responses elicited by variant-updated vaccines. It may be more important to reproduce exposures with the ancestral strain and an Omicron lineage as a means to approximate the immune experience of the general population.

Despite the importance of immune-experienced models, it is critical to also evaluate vaccine immunogenicity in naïve models that are particularly relevant to younger age cohorts that have not yet been exposed to SARS-CoV-2 through either infection or vaccination. Children under the age of four experience a high burden of COVID-19 hospitalization. During the 2023–2024 respiratory season in the US, the cumulative COVID-19 hospitalization rate for children under four was 104.3 per 100,000, and was higher only among persons ≥50 years of age ^4^. Notably, 50% of infants, children, and adolescents ≤17 years hospitalized with COVID-19 from July 2023–March 2024, and 40% of those admitted to the ICU, had no underlying medical conditions ^49^. According to 2023 data in the US, COVID-19 was the third leading cause of death due to infectious disease in individuals 0–17 years of age ^50^. Given this public health relevance, we evaluated a two-dose primary series in naïve mice, wherein we found that both JN.1 and KP.2-adapted vaccines elicited a similar breadth of neutralizing activity against all JN.1 sublineages that were improved over the XBB.1.5 vaccine responses by an order of magnitude (8-to-29-fold higher). Responses elicited by the XBB.1.5 vaccine were also higher (10-fold) than the prior vaccine (bivalent WT + Omicron BA.4/5) in a previous study ^23^. Although the comparative trends in neutralizing responses across vaccine groups were consistent across vaccine-experienced and naïve mouse studies, the magnitude of those differences were much greater in the naïve model, reflecting the considerable antigenic distance between the XBB and JN.1 lineage clusters.

Overall, JN.1 and KP.2 BNT162b2 vaccines elicited improved neutralizing antibody responses over the XBB.1.5 vaccine in mice against the contemporary JN.1 lineages. Additionally, the maintenance of comparable T cell responses between vaccines across a broad set of variant-specific peptides in mice after booster vaccination indicates that cellular immune responses are not substantially impacted by the mutational differences among recent Omicron lineages. Biophysical characterization indicated that JN.1 and KP.2 S proteins are relatively similar to one another but differ in their structural features from prior Omicron lineages. As XBB.1.5 vaccines have demonstrated diminished effectiveness against the JN.1 lineage cluster, and preclinical data have trended closely with clinical responses and real-world effectiveness outcomes, it is anticipated that the potent responses elicited by both the JN.1 and KP.2-adapted BNT162b2 vaccines will confer robust protection against antigenically drifted JN.1 sublineages, including the most globally dominant and recently rising strains, similar to how the XBB.1.5 vaccine performed against the XBB lineage cluster ^16–18,34^. As reported here, and moving forward into the current and future peak seasons of COVID-19 disease activity, a multidimensional approach is needed to monitor the evolutionary dynamics and epidemiology of novel SARS-CoV-2 lineages and the effectiveness of the JN.1- and KP.2-adapted vaccines in protecting against a wide range of clinical outcomes.

## Acknowledgements

We thank the Pfizer Comparative Medicine department and veterinary staff at Pearl River, NY for their contributions to the *in vivo* studies; members of Pfizer Vaccine Research and Development at Pearl River, NY, for their contributions to assay development and implementation, virus variant monitoring, material production and characterization, and material and technical support in molecular biology, virology and immunology; and members of Pfizer Discovery Sciences in Groton, CT, for their material and technical support on recombinant protein production and characterization. The authors also thank Teresa Hauguel for review of the manuscript, Christina D’Arco for writing and graphics support and Charlotte Ford for editorial assistance.

## Author Contributions

WC contributed to writing of the original draft, methodology, investigation, and data generation and interpretation. KRT contributed to the writing of the original draft, investigation, data generation and interpretation. IWW contributed to writing of the original draft, cryo-EM structural determination and visualization, data analysis and interpretation. LTM, MR, WL, SS, SR, KPD, AY, DS, SM, SS, WS, RMM and PMD contributed to data generation, analysis, visualization and interpretation; LTM also contributed to serologic methodology. JSC performed antigen purification and thermal shift experiments. PS performed BLI experiments. KFF performed recombinant protein expression. TJM and GMW performed site-specific glycan mapping by MS studies. APM contributed to data analysis. CIC and AM contributed to data interpretation. US and ASA supervised the work. HW contributed to writing of the original draft, supervision, investigation, visualization, data interpretation and analysis related to the structural and biophysical work. KAS contributed to writing of the original draft, data interpretation, investigation, and overall supervision and conceptualization of the work. KM contributed to writing of the original draft, experimental designs, data analysis and interpretation, investigation and overall supervision and conceptualization of the work. All authors contributed to the development and critical review of the manuscript and have read and agreed to the published version.

