## Supplementary Materials for "Immunologic and Biophysical Features of the BNT162b2 JN.1- and KP.2-Adapted COVID-19 Vaccines"

#### Supplementary Methods

*SARS-CoV-2 S(P2) and RBD Protein Expression and Purification*

Expi293F cells (Thermo Fisher Scientific) grown in Expi293 medium were transiently transfected with S or RBD protein expression constructs in the pcDNA3.1(+) constitutive expression vector. Expression was conducted at 37 °C for 24 hours before adding Expectation enhancers (Thermo Fisher Scientific). After addition of enhancers, the temperature was dropped to 32° C and expression was allowed for another 48-72 hours before collecting. A modified protocol of procedures described by Zhang et al [1] was used for purification of the SARS-CoV-2 FL S(P2). Briefly, the transfected cells were lysed in a solution containing Buffer A (100 mM HEPES pH 8.0, 150 mM NaCl, 1 mM EDTA), 1% (w/v) *n*-dodecyl-*β*-D-maltopyranoside (DDM, Anatase), EDTA-free complete protease inhibitor cocktail (Roche), and Pierce Universal Nuclease (Thermo Fisher) at 4 °C for 1 h. After a clarifying spin at 40,000 × g for 45 min, the supernatant was filtered with 0.2 mm filter (Nalgene 78018-24, 1 LL) before batch bound onto StrepTactin HP resin (Cytiva) equilibrated with the lysis buffer at 4 °C for 1 h. Resin was collected by centrifugation at 1000 × g and loaded onto EconoColumn (Bio-Rad) for gravity flow purification. The column was washed with Buffer A containing 0.5% DDM, 10 mM ATP, and 10 mM $MgCl_2$, followed by additional washes with Buffer A and gradually reduced concentrations of DDM (0.5% - 0.02%). FL S(P2) was eluted with Buffer A containing 0.02% DDM and 5 mM d-Desthiobiotin. The protein was further purified by size exclusion chromatography (SEC) on a Superose 6 10/300 column (Cytiva) in a buffer containing 25 mM Tris pH 7.5, 150 mM NaCl, 1 mM EDTA, and 0.02% DDM. DDM-purified FL S(P2) was eluted as a single peak over SEC. FL S(P2) protein from the SEC peak fractions were analyzed by denaturing PAGE using a 4-15% Criterion TGX Stain-Free Gel (Bio-Rad), and used in thermostability ($T_m$), biolayer interferometry (BLI), mass spectrometry and cryogenic electron microscopy (cryo-EM) experiments. Affinity-tagged RBDs were expressed in Expi293F cells and purified from cell culture medium at 120h using affinity purification and Superdex200 gel filtration columns (Cytiva). Proteins were stored in 100 mM Tris pH7.5, 150 mM NaCl, and 10% glycerol.

*Cryo-EM Data Processing and Model Building for JN.1 and KP.2 S(P2)*

Cryo-EM data was processed using CryoSPARC version 4.4.1. Movies were patch motion corrected and micrographs with >5Å estimated resolution were rejected following CTF estimation. A subset of 1k JN.1 movies were blob picked and extracted with a 3-fold down sampled pixel size. An initial 4.5Å Nyquist-limited, C3-symetric reconstruction was obtained from two round of 2D classification, ab initio reconstruction, and non-uniform refinement. The full JN.1 dataset was picked using 16 templates generated with the initial map. 2D classification and heterogenous refinement of the JN.1 dataset and non-uniform refinement of full-pixel size particle stacks yielded a C3-symetric reconstruction with all 3 RBD in the down conformation (3-down) and a conformer with one RBD up (1-up). The initial high resolution JN.1 1-up map was used to build a model as described below. This model was used to prepare reference coordinates of 3-down, 1-up, 2-up, and 3 up RBD conformers in PyMOL. The molmap feature of ChimeraX [2] was used to generate 8Å resolution 3D volumes that were used as 3D classes in heterogenous refinement following an initial curation with 2D classification and 3D heterogeneous refinement of template-picked particle stacks for both the JN.1 and KP.2 datasets. The first round of reference-based heterogenous refinement of curated datasets yielded a single good 3-down class. A second round led to good classes for 3-down and 1-up classes for both datasets and an additional 2-up class for KP.2. Particles from each good 3D class were subjected to two additional rounds of heterogenous refinement and a final non-uniform refinement. The final maps were local resolution filtered to improve map completeness for poorly resolved regions of the NTD and RBD.

Model building was conducted by generating models using AlphaFold2 [3] for the JN.1 NTD, RBD, SD1, and SD2 domains and extracting the S2 domain from the PDB entry 6xr8. Template models were rigid body docked in EM densities using UCSF ChimeraX. Conformational fluctuations of the NTD and RBD necessitated docking these domains into each map. Coordinates of the domains and junctions were manually revised and point mutations corrected using Coot [3]. Coordinates were refined using Phenix real-space refinement [4] with secondary structure restraints imposed

*Flow Cytometry Analysis of Human Angiotensin Converting Enzyme (hACE2)-Peptidase Domain (PD) Binding to Cell Surface-Expressed S(P2) Protein*

Expi293F cells were collected 48 hours post-transfection with P2 S proteins. Cells were incubated for one hour at room temperature in TBS + 4% BSA + 0.01 mg/ml 7-AAD to detect non-viable cells and with 20 nM His-tagged hACE2-PD and 13 nM FITC-labeled anti-His to detect FL S surface expression. The incubation was done in Corning V-bottom 96-well plates (Corning, cat # 3897) followed by TBS wash before the plate was read by Guava EasyCyte HT flow cytometer. For each condition, 3 replicates were measured with 5000 events collected per replicate.

*BNT162b2 Variant-Adapted Vaccine Formulations*

The monovalent XBB.1.5-adapted BNT162b2 vaccine used in this study was previously described [5]. The monovalent JN.1-adapted and monovalent KP.2-adapted BNT162b2 vaccines encode S(P2) with changes from the original BNT162b2 limited to variant-specific S sequence changes in Omicron JN.1 (EPI_ISL_18374006) and KP.2 variants (EPI_ISL_18945750). Purified nucleoside-modified RNA was formulated into lipid nanoparticles as previously described [6].

*Animal Blood Collection and Splenocyte Isolation*

For interim and terminal bleeds, blood was collected as previously described [5]. For flow cytometry of murine splenocytes, spleens were collected from five mice per group at the terminal time point for each study. Spleens were harvested and processed as described previously [5]. At each study end, mice were euthanized under a surgical plane of isoflurane and cervical dislocation was performed as a secondary method to confirm death.

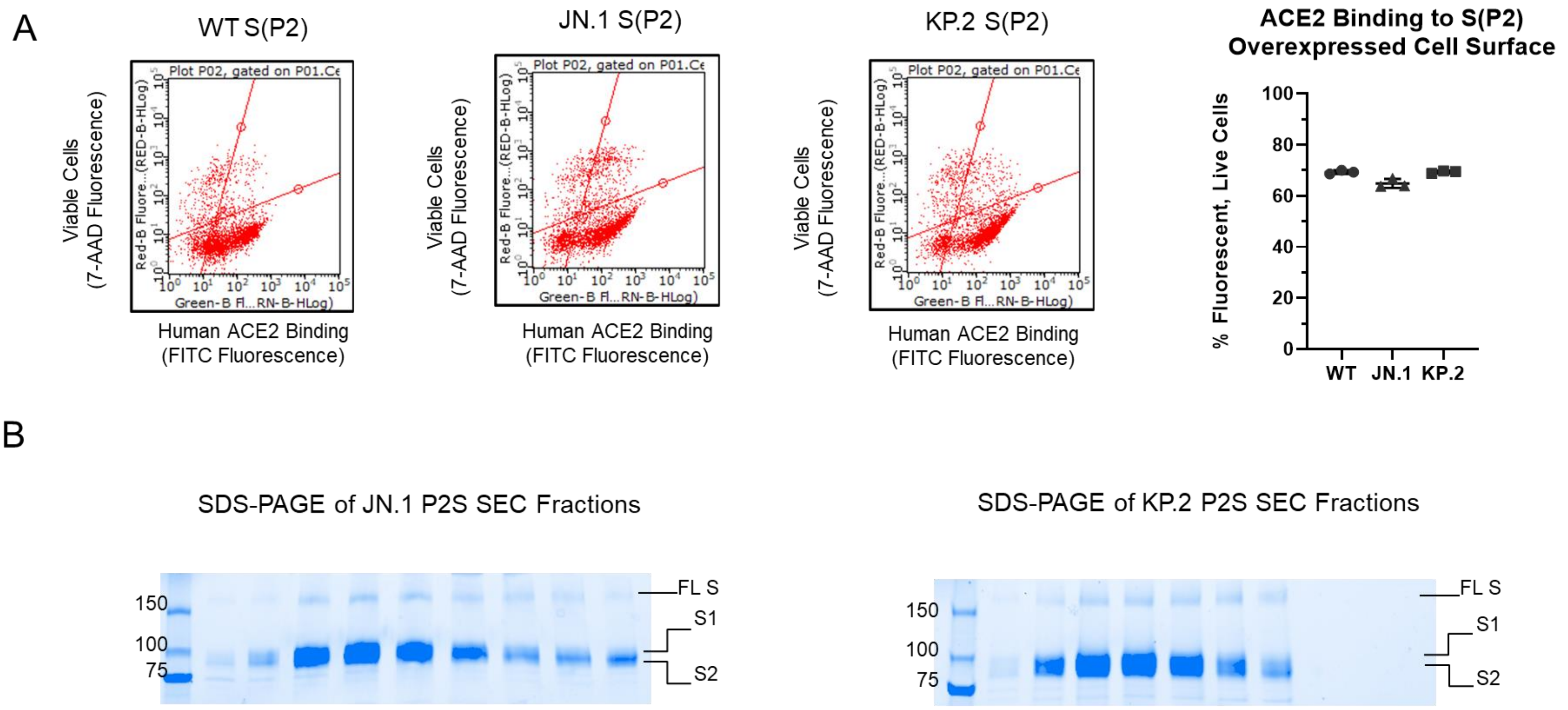

**Figure S1. Additional Biophysical Data of Recombinantly Expressed S(P2) Proteins. a** ACE2-PD binding to cell surface-expressed S(P2) proteins. Flow cytometry experiments were collected with Expi293F cells overexpressing S(P2) under nonpermeabilized conditions. The nucleic acid dye 7-AAD was used to differentiate live and dead cells (y axis). S(P2) expression was confirmed by staining with ACE2-PD complexed by a FITC-labeled anti-His antibody (x axis). The lower right quadrants in the flow plots indicate the live cell population that has functional S(P2) expressed to the cell surface. The percentages of live cells with detectable S(P2) expression on the cell surface from three replicates ($n = 3$) are quantified and shown. **b** SDS-PAGE of DDM-solubilized FL S(P2) after size exclusion chromatography (SEC) purification.

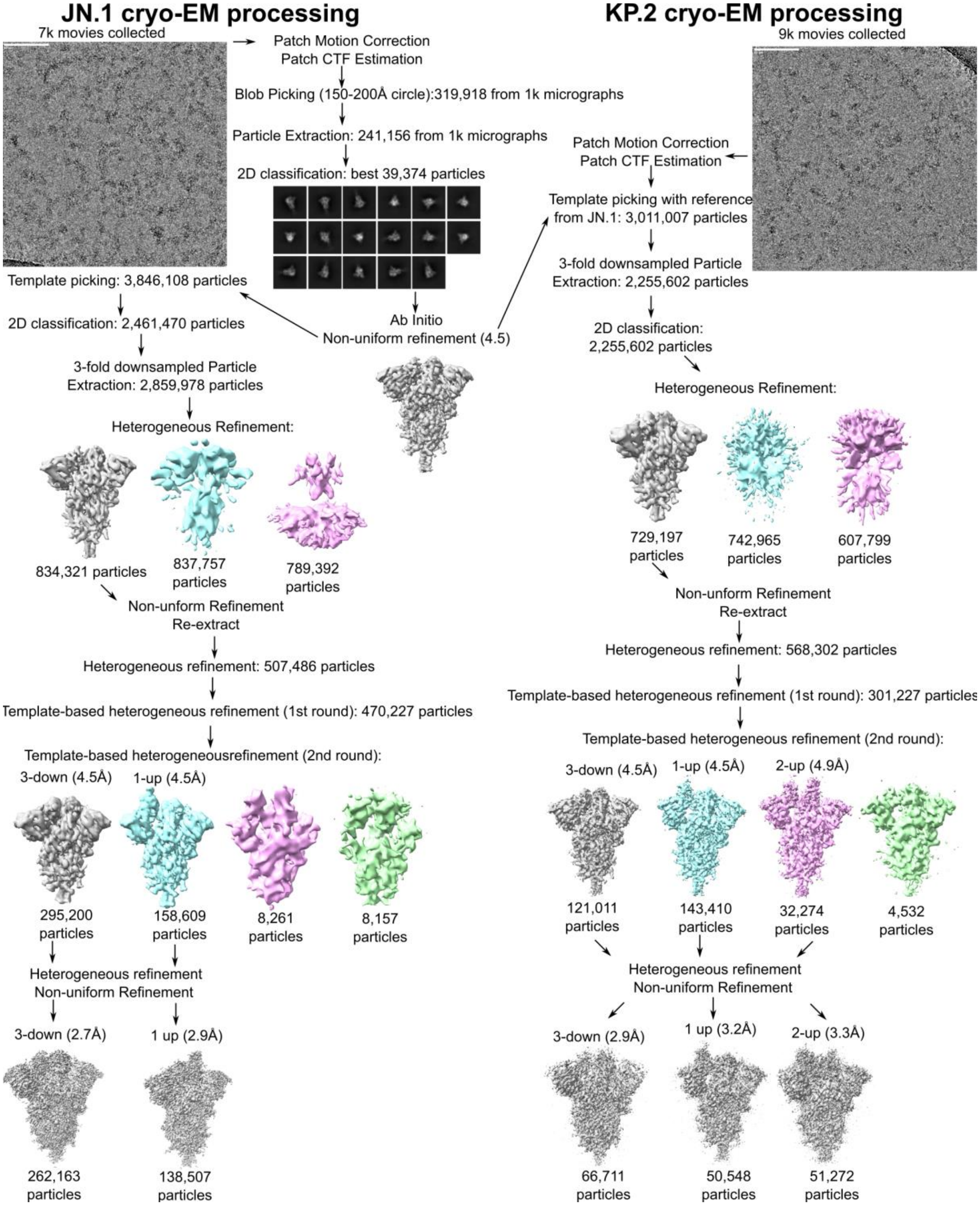

**Figure S2. Cryo-EM Data Processing.** Key decisions are depicted for the processing of the JN.1 (left) and KP.2 (right) datasets. Heterogeneous refinement was employed to both reject bad particles and identify the relevant RBD conformational states. Many particles were removed in the final round of heterogenous refinement of the KP.2 conformations to obtain high-resolution reconstructions so the particles from the 2$^{nd}$ round of heterogenous refinement are used to calculate the proportions in Figure 2.

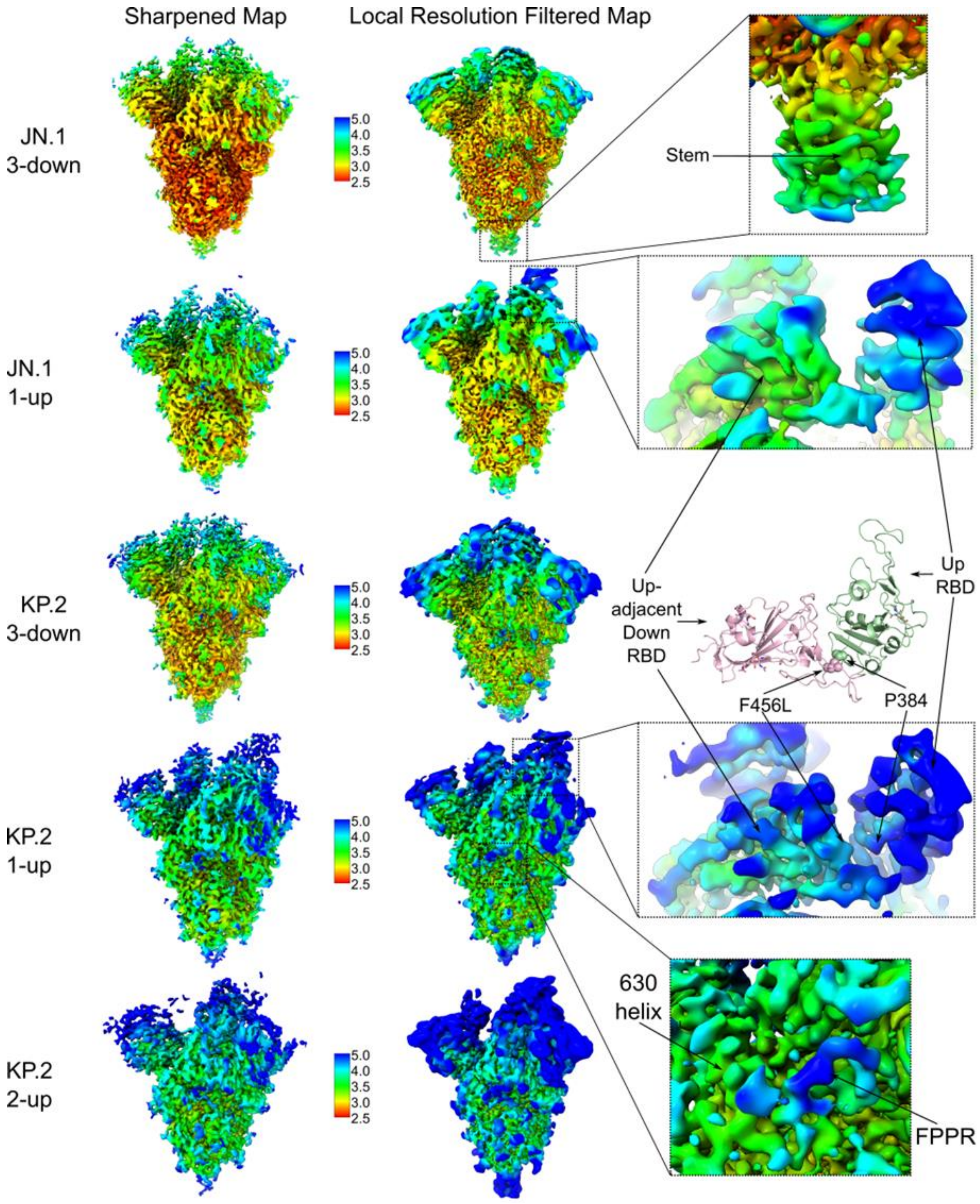

**Figure S3. Local Resolution Analysis of Cryo-EM Densities.** Estimated local resolutions were mapped on to the final sharpened 3D reconstructions (left) and utilized to generate and color local resolution filtered maps (middle). Local resolution filtered maps were used for model building and also shown in Fig. 2. Close up views of key features highlighted in the main text and Figure 2 are shown on the right.

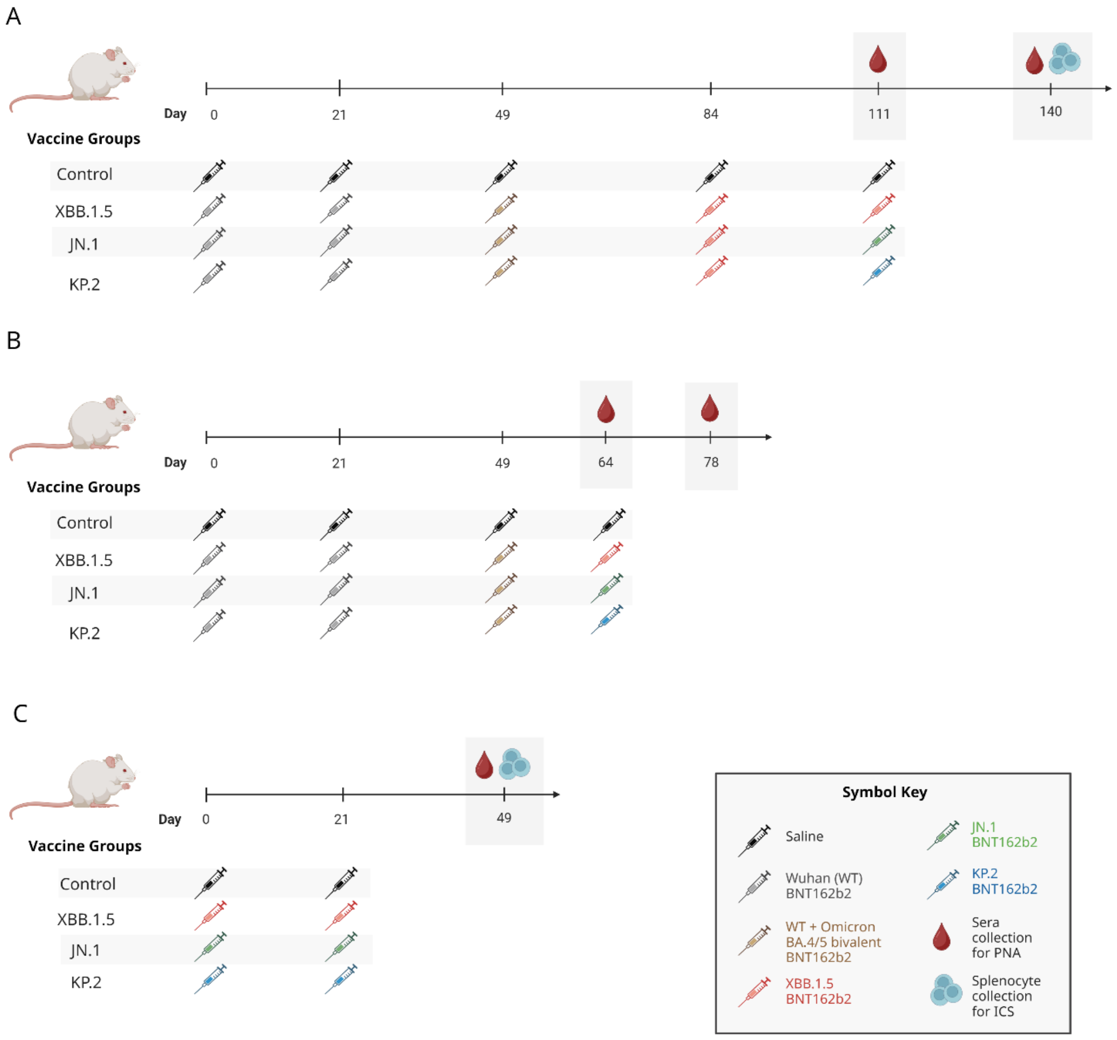

**Figure S4. BNT162b2 XBB.1.5, JN.1 and KP.2 Mouse Immunogenicity Study Designs. a-c** Schematics are shown for (**a** and **b**) two booster immunogenicity studies and (**c**) one primary series study of monovalent BNT162b2 XBB.1.5, JN.1 and KP.2 vaccines conducted in female BALB/c mice. **a** mice were vaccinated intramuscularly (i.m.) with 2 doses (D0, D21) of original BNT162b2 WT vaccine, followed by a 3rd dose booster (D49) of bivalent WT + Omicron BA.4/5 vaccine, a 4th dose booster (D84) of the monovalent XBB.1.5-adapted vaccine, and a 5th booster (D111) of the monovalent XBB.1.5, JN.1, or KP.2-adapted vaccine. Sera were collected twice for evaluation (D111, D140) in a pseudovirus neutralization assay (PNA) and spleens were collected at the study end (Day 140) to evaluate cell-mediated immune responses in flow cytometry-based intracellular cytokine staining (ICS). **b** mice were vaccinated i.m. according to the schedule shown in (A), except that the 4th and final booster dose (D64) was the monovalent XBB.1.5, JN.1, or KP.2-adapted vaccine. Sera were collected twice (D64, D78) for evaluation in a PNA. **c** mice were vaccinated i.m. with 2 doses (D0, D21) of the monovalent XBB.1.5, JN.1, or KP.2-adapted vaccine. Sera were collected for evaluation in a PNA and spleens were collected (5 mice/group) on D49 for evaluation in ICS. For A, B, and C, mice were vaccinated in groups of 10 and a control group (10 mice) received saline injections according to the active vaccine group schedule. The total dose level for each vaccine formulation was 0.5 µg. Created in BioRender. D'Arco, C. (2024) BioRender.com/g65e540

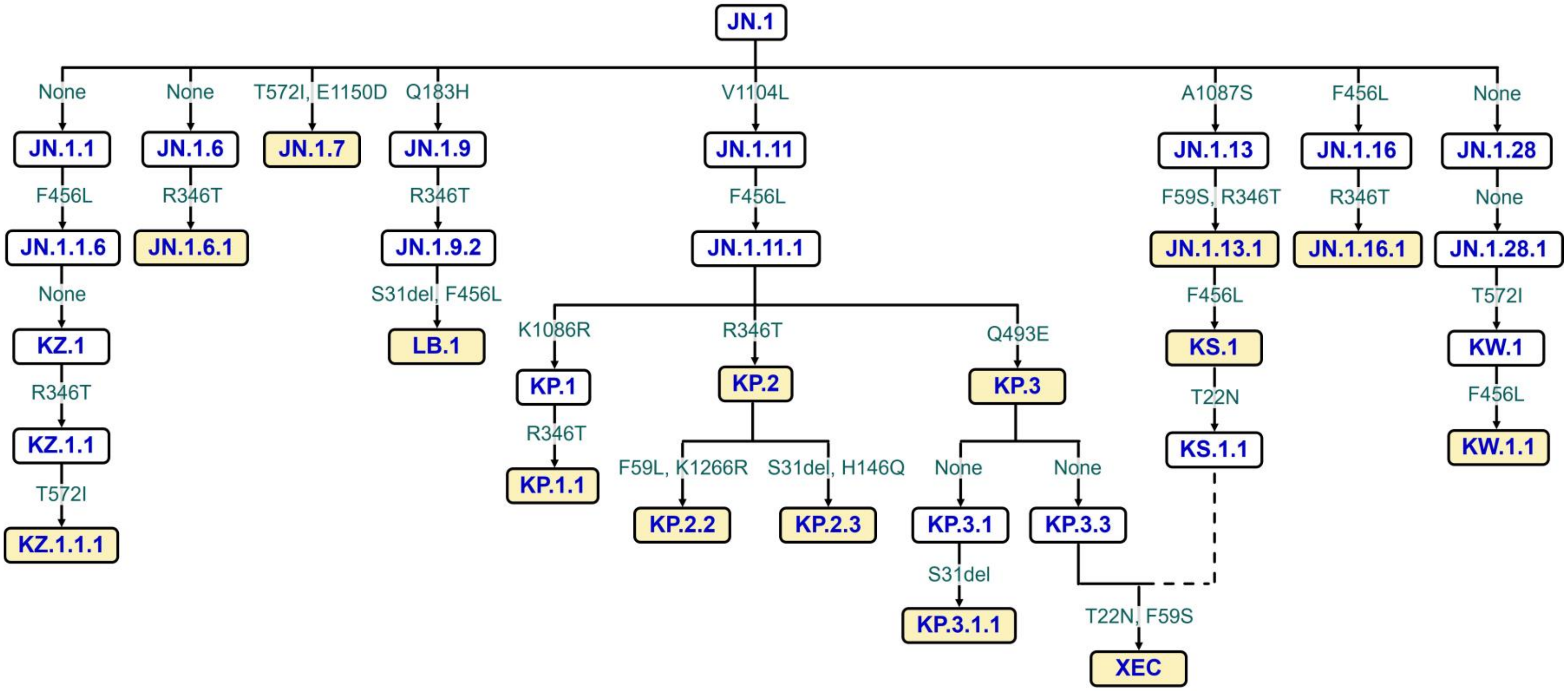

**Figure S5. Acquired Amino Acid Substitutions in the Spike Protein of SARS-CoV-2 Lineages Evaluated in a Pseudovirus Neutralization Assay.** This tree highlights the key amino acid changes in the SARS-CoV-2 spike protein (displayed in green font) of lineages descended from the JN.1 variant. JN.1 and descendant lineages that were deemed most epidemiologically and immunologically relevant (shown in yellow boxes) were included in the pseudovirus panel for testing vaccine-elicited neutralizing activity.

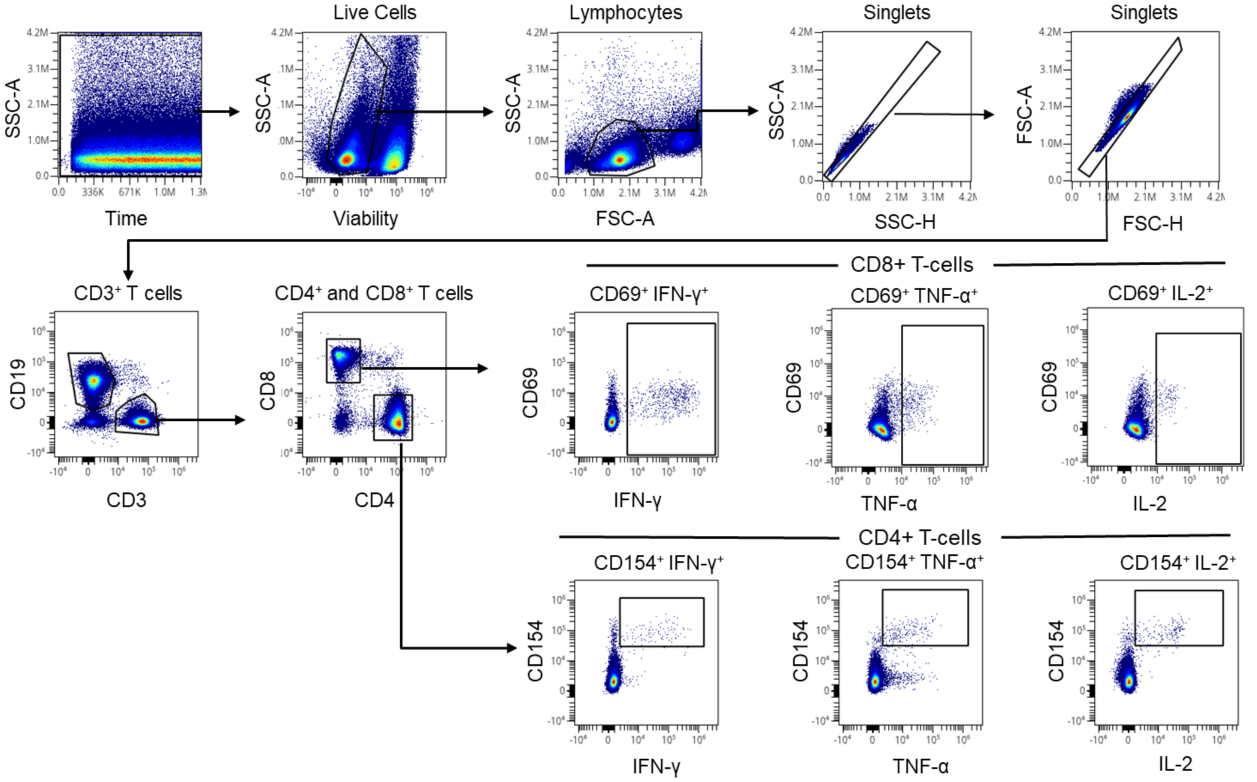

**Figure S6. Gating Strategy for Intracellular Cytokine Staining Flow Cytometry Analysis of T Cell Responses.** Flow cytometry gating strategy for identification of SARS-CoV-2 spike-specific T cells for different sublineages. (Upper row, left to right) Starting with events acquired with a constant flow stream and fluorescence intensity, viable cells, lymphocytes, and single events were identified and gated. Within singlet lymphocytes, CD19-negative CD3+ T cells were identified and gated into $CD4^+$ and $CD8^+$ T cells (middle row). Activated $CD8^+$ T cells were identified by gating on CD69 and cytokine-positive cells (middle row). Antigen-specific $CD4^+$ T cells were identified by gating on CD154 and cytokine-positive cells (bottom row). The antigen-specific T cell frequencies were used for further analysis**.**

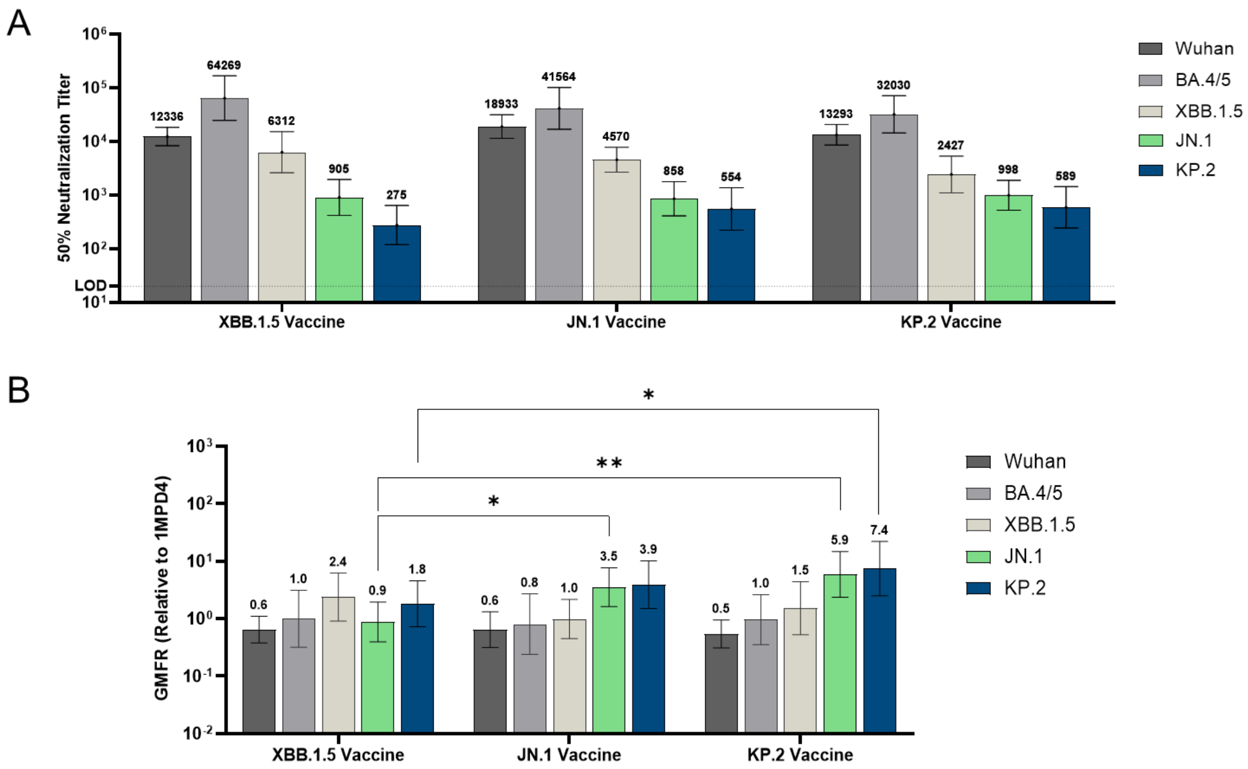

**Figure S7. Geometric Mean Fold Rise in Pseudovirus Neutralization Titers ($NT_{50}$) from Pre- to Post-5th Dose with BNT162b2 JN.1 and KP.2-Adapted Vaccines in BNT162b2-Experienced Mice.** Female BALB/c mice (10/group) vaccinated as shown in Fig. S5A with two-doses of original monovalent BNT162b2 (WT) vaccine, one subsequent dose of bivalent WT + Omicron BA.4/5 vaccine, and one subsequent dose of the monovalent XBB.1.5-adapted vaccine, received a single intramuscular booster dose of one of the following variant-modified BNT162b2 vaccines: monovalent XBB.1.5, JN.1, or KP.2. Serum neutralizing antibody responses were measured by a pseudovirus neutralization assay (PNA). **a** 50% pseudovirus neutralization titers are shown as geometric mean titers (GMT) with 95% CI of 10 mice per vaccine group on Day 111, pre-5$^{th}$ dose. The limit of detection (LOD) is the lowest serum dilution, 1:20. **b** The fold rise in geometric mean neutralizing titers (GMFR) from pre-fifth dose to one-month post-fifth dose are shown above each bar for the Wuhan (WT) reference strain and Omicron BA.4/5, XBB.1.5, JN.1 and KP.2 variants with 95% CI. The LOD is the lowest serum dilution, 1:20. Asterisks indicate statistical significance of pseudovirus GMFR comparisons between vaccine groups. ** $p<0.01$, * $p<0.05$.

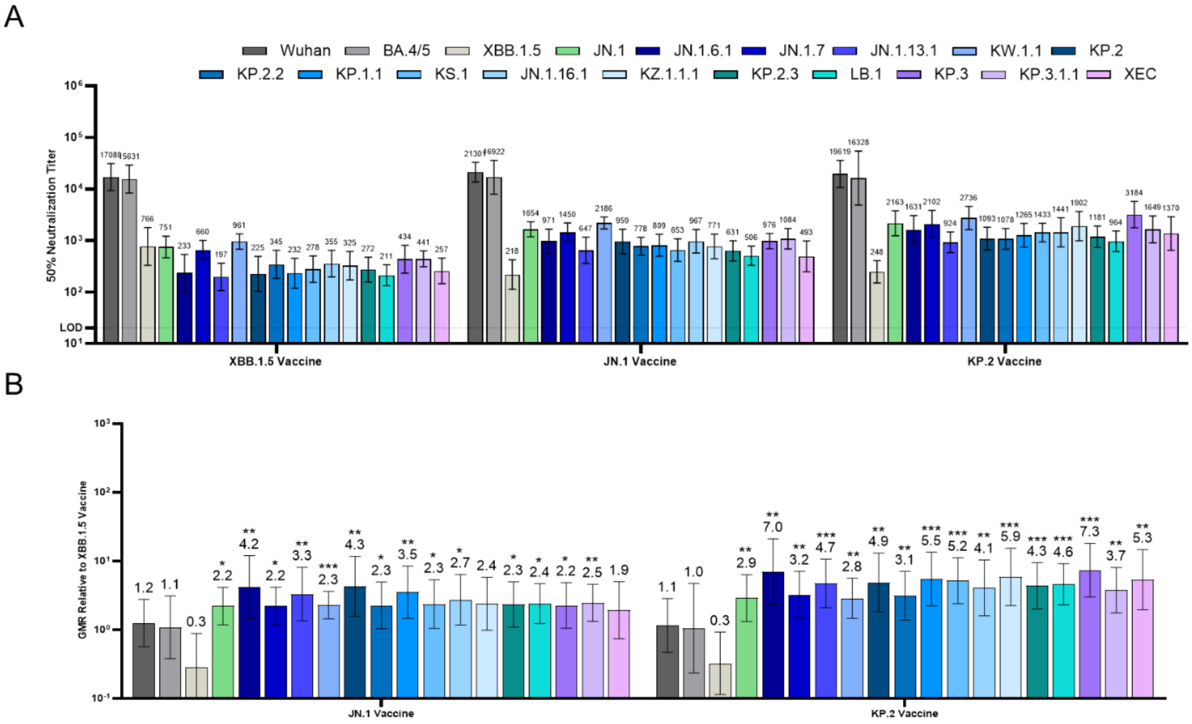

**Figure S8. Pseudovirus Neutralization Elicited by BNT162b2 XBB.1.5, JN.1 and KP.2-Adapted Vaccines Administered to BNT162b2-Experienced Mice as a 4th Dose.** Female mice were immunized i.m. according to Fig. S5B on Days 0 and 21 with the BNT162b2 Wuhan (WT), on Day 49 with the bivalent BNT162b2 (WT + BA.4/5), and on Day 64 with either the BNT162b2 XBB.1.5, JN.1 or KP.2 vaccine. Two weeks post-fourth dose, sera were collected from the terminal bleed and neutralizing antibody responses against a panel of 19 pseudoviruses were measured that included the Wuhan (WT) reference strain and Omicron variants BA.4/5, XBB.1.5, JN.1, JN.1.6.1, JN.1.7, JN.1.13.1, KW.1.1, KP.2, KP.2.2, KP.1.1, KS.1, JN.16.1, KZ.1.1.1, KP.2.3, LB.1, KP.3, KP.3.1.1, and XEC. **a** The number above each bar indicates the 50% neutralizing geometric mean titer (GMT) with 95% CI of 10 mice per vaccine group. **b** The geometric mean ratio (GMR) is shown as the ratio of the BNT162b2 KP.2 or JN.1 vaccine GMT to the BNT162b2 XBB.1.5 vaccine GMT of the corresponding pseudovirus. The number above each bar indicates the GMR with 95% CI. The limit of detection (LOD) is the lowest serum dilution, 1:20. Asterisks indicate statistical significance of pseudovirus GMR relative to the corresponding pseudovirus in the monovalent XBB.1.5 vaccine group. ***$p<0.001$, ** $p<0.01$, * $p<0.05$

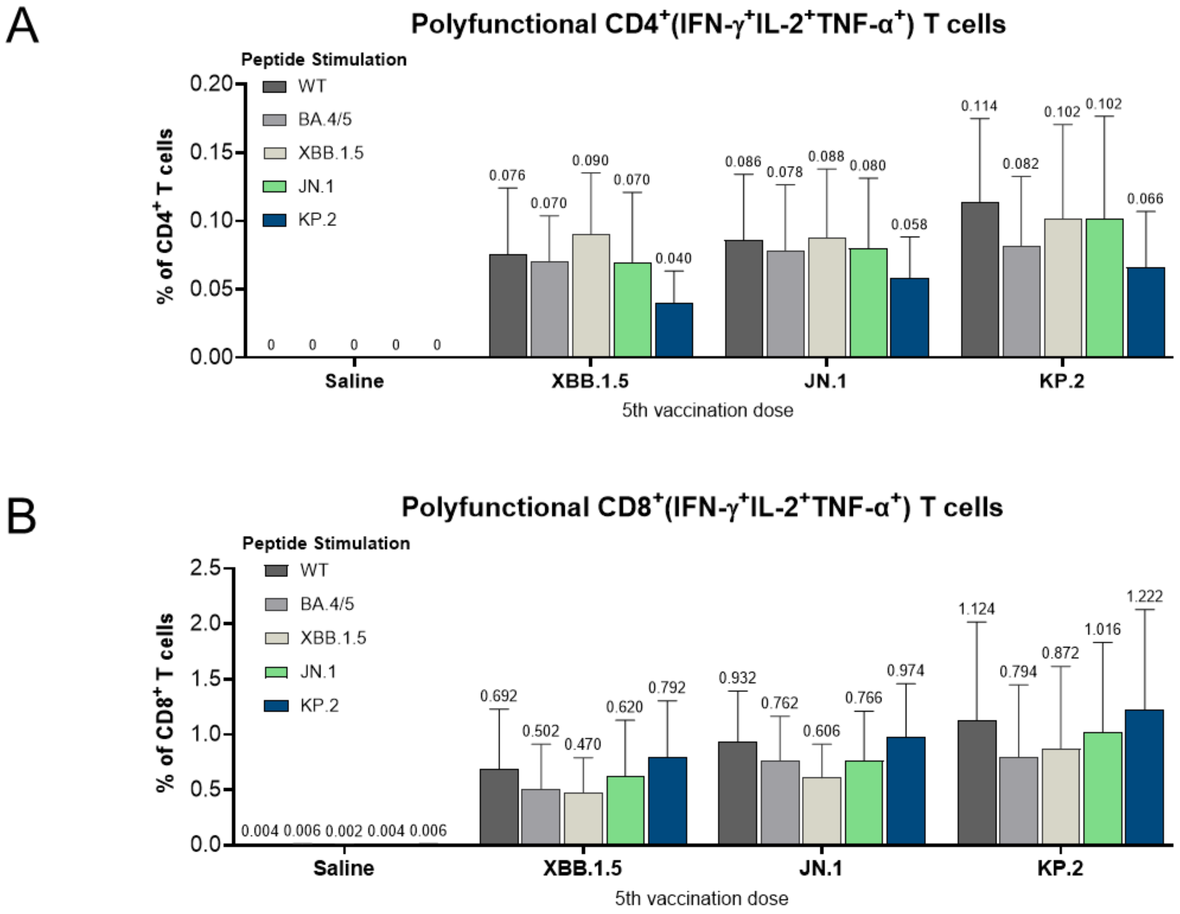

**Figure S9. Polyfunctional T Cells Elicited by BNT162b2 XBB.1.5, JN.1 and KP.2-Adapted Vaccines Administered as a 5th Dose to BNT162b2-Experienced Mice.** One-month after the fifth dose of BNT162b2 variant-adapted vaccine (XBB.1.5, JN.1, or KP.2) or saline (negative control), S-specific splenocytes (n=5/group) were characterized by a flow cytometry-based intracellular cytokine staining (ICS) assay. All samples were stimulated *ex vivo* with S peptide pools from the WT reference strain and Omicron BA.4/5, XBB.1.5, JN.1, and KP.2 sublineages. **a-b** Graphs show the frequency of (**a**) polyfunctional $CD4^+$ T cells (IFN-$\gamma^+$ TNF-$\alpha^+$ IL-$2^+$) and (**b**) polyfunctional $CD8^+$ T cells (IFN-$\gamma^+$ TNF-$\alpha^+$ IL-$2^+$) in response to stimulation with each peptide pool across vaccine groups. Bars depict mean frequency + SEM.

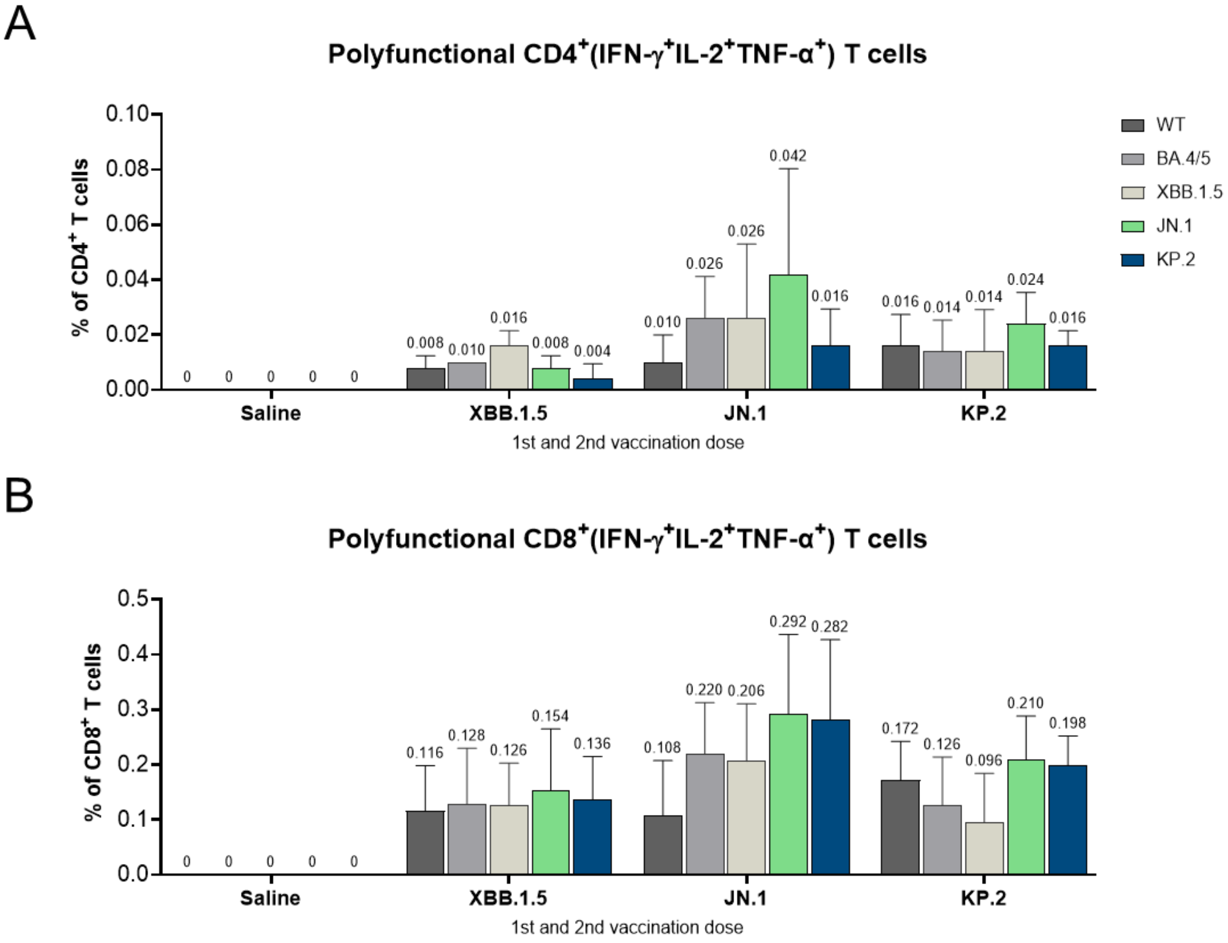

**Figure S10. Polyfunctional T Cells Elicited by BNT162b2 XBB.1.5, JN.1 and KP.2-Adapted Vaccines Administered to Naïve Mice.** At one-month post-second dose (completion of primary series) of BNT162b2 variant-adapted vaccine (XBB.1.5, JN.1, or KP.2) or saline (negative control), S-specific splenocytes (n=5/group) were characterized by a flow cytometry-based intracellular cytokine staining (ICS) assay. All samples were stimulated ex vivo with S peptide pools from the WT reference strain and Omicron BA.4/5, XBB.1.5, JN.1, and KP.2 sublineages. **a-b** Graphs show the frequency of **(a)** polyfunctional $CD4^+$ T cells (IFN-$\gamma^+$ TNF-$\alpha^+$ IL-$2^+$) and **(b)** polyfunctional $CD8^+$ T cells (IFN-$\gamma^+$ TNF-$\alpha^+$ IL-$2^+$) in response to stimulation with each peptide pool across vaccine groups. Bars depict mean frequency + SEM.

**Table S1. Expression Constructs of FL S Proteins**

| Description | Proline Mutations | Spike Mutations | Affinity Tag |
|---|---|---|---|
| P2 S (Original Variant) | K986P, V987P | | C-terminal TwinStrep |
| P2 S (JN.1) | K986P, V987P | ins16MPLF, T19I, R21T, L24del, P25del, P26del, A27S, S50L, H69del, V70del, V127F, G142D, Y144del, F157S, R158G, N211del, L212I, V213G, L216F, H245N, A264D, I332V, G339H, K356T, S371F, S373P, S375F, T376A, R403K, D405N, R408S, K417N, N440K, V445H, G446S, N450D, L452W, L455S, N460K, S477N, T478K, N481K, V483del, E484K, F486P, Q498R, N501Y, Y505H, E554K, A570V, D614G, P621S, H655Y, N679K, P681R, N764K, D796Y, S939F, Q954H, N969K, P1143L | C-terminal TwinStrep |
| P2 S (KP.2) | K986P, V987P | ins16MPLF, T19I, R21T, L24del, P25del, P26del, A27S, S50L, H69del, V70del, V127F, G142D, Y144del, F157S, R158G, N211del, L212I, V213G, L216F, H245N, A264D, I332V, G339H, R346T, K356T, S371F, S373P, S375F, T376A, R403K, D405N, R408S, K417N, N440K, V445H, G446S, N450D, L452W, L455S, F456L, N460K, S477N, T478K, N481K, V483del, E484K, F486P, Q498R, N501Y, Y505H, E554K, A570V, D614G, P621S, H655Y, N679K, P681R, N764K, D796Y, S939F, Q954H, N969K, V1104L, P1143L | C-terminal TwinStrep |
| RBD (Original Variant) | N/A | | C-terminal StrepII |
| RBD (JN.1) | N/A | I332V, G339H, K356T, S371F, S373P, S375F, T376A, R403K, D405N, R408S, K417N, N440K, V445H, G446S, N450D, L452W, L455S, N460K, S477N, T478K, N481K, V483del, E484K, F486P, Q498R, N501Y, Y505H | C-terminal His×8 |
| RBD (KP.2) | N/A | I332V, G339H, R346T, K356T, S371F, S373P, S375F, T376A, R403K, D405N, R408S, K417N, N440K, V445H, G446S, N450D, L452W, L455S, F456L, N460K, S477N, T478K, N481K, V483del, E484K, F486P, Q498R, N501Y, Y505H | C-terminal His×8 |

### Table S2. Glycosylation Heterogeneity of JN.1 and KP.2 Spike Proteins

| CoVid JN.1 pSB7938 | | | | |
|---|---|---|---|---|
| **Site** | **Glycan** | **Potential Glycan Structures** | **Number Detected** | **Ion Intensity** |
| n(245)RS | | | | 6.41E+07 |
|  | HexNAc(1) |  | 1 | 5.71E+07 |
|  | HexNAc(2)Hex(1) |  | 1 | 8.65E+06 |
|  | HexNAc(2)Hex(2) |  | 1 | 7.93E+06 |
| | HexNAc(2)Hex(3) | 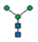 | 1 | 6.72E+07 |
| | HexNAc(2)Hex(3)Fuc(1) | 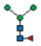 | 1 | 8.44E+06 |
| | HexNAc(2)Hex(4) | 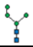 | 1 | 2.60E+08 |
| | HexNAc(2)Hex(5) | 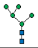 | 1 | 6.34E+09 |
| | HexNAc(2)Hex(6) | 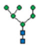 | 1 | 2.98E+09 |
| | HexNAc(2)Hex(7) | 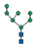 | 1 | 2.14E+09 |
| | HexNAc(2)Hex(8) | 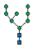 | 1 | 1.91E+09 |
| | HexNAc(2)Hex(9) | 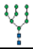 | 1 | 8.52E+08 |
| | HexNAc(3)Hex(3) | 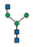 | 1 | 2.88E+07 |
| | HexNAc(3)Hex(3)Fuc(1) | 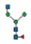 | 1 | 1.75E+07 |
| | HexNAc(3)Hex(4) | 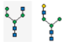 | 1 | 2.91E+07 |
| | HexNAc(3)Hex(4)Fuc(1) | 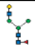 | 1 | 2.58E+07 |
| | HexNAc(3)Hex(4)NeuAc(1) | 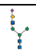 | 1 | 1.45E+07 |
| | HexNAc(3)Hex(4)NeuAc(1)Fuc(1) | 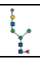 | 1 | 4.21E+06 |
| | HexNAc(3)Hex(5) | 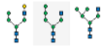 | 1 | 7.25E+07 |
| | HexNAc(3)Hex(6) | 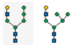 | 1 | 8.47E+07 |
| | HexNAc(4)Hex(3) | 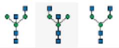 | 1 | 4.88E+06 |
| | HexNAc(4)Hex(3)Fuc(1) | 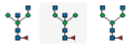 | 1 | 1.65E+07 |
| | HexNAc(4)Hex(4)Fuc(1) | 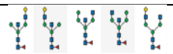 | 1 | 3.58E+06 |
| | HexNAc(4)Hex(4)NeuAc(1)Fuc(1) | 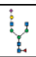 | 1 | 3.26E+06 |
| | HexNAc(4)Hex(5) | 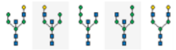 | 1 | 6.44E+06 |
| | HexNAc(4)Hex(5)Fuc(1) | 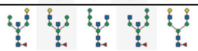 | 1 | 1.67E+07 |
| | HexNAc(4)Hex(5)Fuc(1)NeuAc(1) | 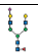 | 1 | 1.73E+07 |
| | HexNAc(4)Hex(5)Fuc(1)NeuAc(2) | 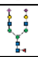 | 1 | 4.62E+06 |

* Ranking was determined by using the summed area extracted ion chromatograms for all the detected charge states and forms for a given glycan using the peptide used to determine the glycosylation level used in the N-glycan level determination.

### CoVid JN.1 pSB7938

| Site | Glycan | Potential Glycan Structures | Number Detected | Ion Intensity |
|---|---|---|---|---|
| n(354)RT | | | 1 | 2.04E+07 |
|  | HexNAc(1) |  | 1 | 1.56E+06 |
| | HexNAc(2)Hex(3) | 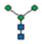 | 1 | 3.35E+06 |
| | HexNAc(2)Hex(4) | 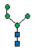 | 1 | 1.87E+07 |
| | HexNAc(2)Hex(5) | 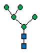 | 1 | 2.45E+08 |
| | HexNAc(2)Hex(6) | 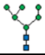 | 1 | 2.81E+08 |
| | HexNAc(2)Hex(7) | 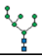 | 1 | 8.94E+08 |
| | HexNAc(2)Hex(8) | 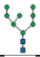 | 1 | 2.66E+09 |
| | HexNAc(2)Hex(9) | 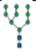 | 1 | 2.62E+09 |
| | HexNAc(3)Hex(3) | 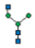 | 1 | 3.92E+06 |
| | HexNAc(3)Hex(4) | 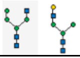 | 1 | 1.18E+07 |
| | HexNAc(3)Hex(4)Fuc(1) | 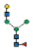 | 1 | 2.93E+06 |
| | HexNAc(3)Hex(4)NeuAc(1) | 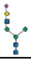 | 1 | 7.54E+05 |
| | HexNAc(3)Hex(5) | 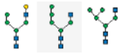 | 1 | 1.59E+07 |
| | HexNAc(3)Hex(6) | 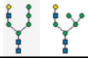 | 1 | 6.85E+06 |
| | HexNAc(4)Hex(3) | 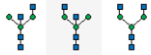 | 1 | 5.68E+06 |
| | HexNAc(4)Hex(3)Fuc(1) | 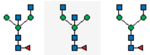 | 1 | 7.38E+06 |
| | HexNAc(4)Hex(4) | 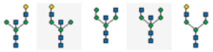 | 1 | 4.66E+06 |
| | HexNAc(4)Hex(4)Fuc(1) | 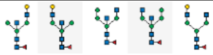 | 1 | 1.40E+07 |
| | HexNAc(4)Hex(4)NeuAc(1) | 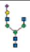 | 1 | 8.90E+05 |
| | HexNAc(4)Hex(4)NeuAc(1)Fuc(1) | 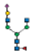 | 1 | 4.97E+05 |
| | HexNAc(4)Hex(5) | 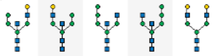 | 1 | 8.69E+06 |
| | HexNAc(4)Hex(5)Fuc(1) | 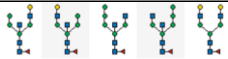 | 1 | 8.77E+06 |
| | HexNAc(4)Hex(5)Fuc(1)NeuAc(1) | 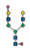 | 1 | 6.09E+06 |
| | HexNAc(5)Hex(4) | 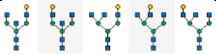 | 1 | 3.70E+06 |
| | HexNAc(5)Hex(4)Fuc(1) | 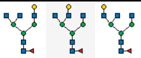 | 2 | 1.04E+07 |

* Ranking was determined by using the summed area extracted ion chromatograms for all the detected charge states and forms for a given glycan using the peptide used to determine the glycosylation level used in the N-glycan level determination.

### CoVid KP.2 pSB8076

| Site | Glycan | Potential Glycan Structures | Number Detected | Ion Intensity |
|---|---|---|---|---|
| n(245)RS | | | 1 | 5.12E+07 |
|  | HexNAc(1) |  | 1 | 3.15E+07 |
| | HexNAc(2)Hex(3) | 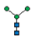 | 1 | 3.56E+07 |
| | HexNAc(2)Hex(3)Fuc(1) | 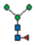 | 1 | 9.01E+06 |
| | HexNAc(2)Hex(4) | 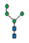 | 1 | 1.60E+08 |
| | HexNAc(2)Hex(5) | 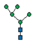 | 1 | 4.06E+09 |
| | HexNAc(2)Hex(6) | 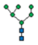 | 1 | 2.06E+09 |
| | HexNAc(2)Hex(7) | 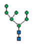 | 1 | 7.95E+08 |
| | HexNAc(2)Hex(8) | 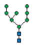 | 1 | 7.18E+08 |
| | HexNAc(2)Hex(9) | 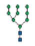 | 1 | 4.01E+08 |
| | HexNAc(3)Hex(3) | 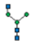 | 1 | 6.90E+07 |
| | HexNAc(3)Hex(3)Fuc(1) | 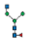 | 1 | 2.51E+07 |
| | HexNAc(3)Hex(4) | 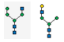 | 1 | 2.00E+07 |
| | HexNAc(3)Hex(4)Fuc(1) | 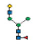 | 1 | 2.40E+07 |
| | HexNAc(3)Hex(4)NeuAc(1) | 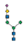 | 1 | 2.74E+07 |
| | HexNAc(3)Hex(4)NeuAc(1)Fuc(1) | 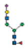 | 1 | 1.12E+07 |
| | HexNAc(3)Hex(5) | 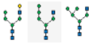 | 1 | 8.32E+07 |
| | HexNAc(3)Hex(6) | 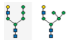 | 1 | 1.13E+08 |
| | HexNAc(4)Hex(3) | 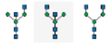 | 1 | 8.66E+06 |
| | HexNAc(4)Hex(3)Fuc(1) | 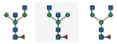 | 1 | 1.72E+07 |
| | HexNAc(4)Hex(4)Fuc(1) | 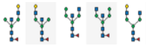 | 1 | 1.45E+06 |
| | HexNAc(4)Hex(4)NeuAc(1)Fuc(1) | 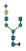 | 1 | 2.40E+06 |
| | HexNAc(4)Hex(5) | 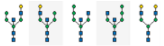 | 1 | 4.67E+06 |
| | HexNAc(4)Hex(5)Fuc(1) | 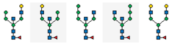 | 1 | 1.74E+07 |
| | HexNAc(4)Hex(5)Fuc(1)NeuAc(1) | 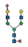 | 1 | 7.22E+06 |
| | HexNAc(4)Hex(5)Fuc(1)NeuAc(2) | 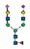 | 1 | 9.26E+06 |
| | HexNAc(4)Hex(6)Fuc(1) | 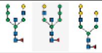 | 1 | Detected |
| | HexNAc(5)Hex(4)Fuc(1) | 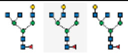 | 1 | 4.27E+06 |
| | HexNAc(5)Hex(5)Fuc(1) | 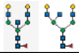 | 1 | 1.77E+06 |

* Ranking was determined by using the summed area extracted ion chromatograms for all the detected charge states and forms for a given glycan using the peptide used to determine the glycosylation level used in the N-glycan level determination.

**Table S3. Cryo-EM Data Collection, Refinement, and Validation Statistics**

| Strain and conformation | JN.1 3-down | JN.1 1-up | KP.2 3-down | KP.2 1-up | KP.2 2-up |
|---|---|---|---|---|---|
| EMDB | 46637 | 46638 | 46639 | 46640 | 46641 |
| PDB | 9D8H | 9D8I | 9D8J | 9D8K | 9D8L |
| **Data collection and processing** | | | | | |
| Magnification | | 165,000 | | 165,000 | |
| Voltage (kV) | | 300 | | 300 | |
| Electron exposure (e–/Å$^2$) | | 40 | | 40 | |
| Defocus range (μm) | | -0.4 to -2.2 | | -0.4 to -2.2 | |
| Pixel size (Å) | | 0.733 | | 0.733 | |
| Initial particle images (no.) | | 3,846,108 | | 3,011,007 | |
| Final particle images (no.) | 262,163 | 138,507 | 66,711 | 50,548 | 54,272 |
| Symmetry imposed | C3 | C1 | C3 | C1 | C1 |
| Map resolution (Å) | 2.7 | 2.9 | 2.9 | 3.2 | 3.3 |
| FSC threshold | 0.143 | 0.143 | 0.143 | 0.143 | 0.143 |
| **Refinement** | | | | | |
| Model resolution (Å) | 2.8 | 3.0 | 2.9 | 3.2 | 3.4 |
| FSC threshold | 0.5 | 0.5 | 0.5 | 0.5 | 0.5 |
| Map sharpening *B* factor (Å$^2$) | -107.5 | -87.4 | -92.5 | -73.1 | -71.6 |
| Model composition | | | | | |
| Non-hydrogen atoms | 27,678 | 27,971 | 27,666 | 27,761 | 27,659 |
| Protein residues | 3,336 | 3,352 | 3,336 | 3,346 | 3,338 |
| Glycans | 63 | 76 | 66 | 67 | 64 |
| *B* factors (Å$^2$) | | | | | |
| Protein | 39.4 | 98.3 | 91.2 | 79.1 | 136.6 |
| Glycan | 76.0 | 123.0 | 117.6 | 118.7 | 168.3 |
| R.m.s. deviations | | | | | |
| Bond lengths (Å) | 0.004 | 0.004 | 0.005 | 0.005 | 0.004 |
| Bond angles (°) | 0.959 | 0.971 | 1.008 | 0.967 | 0.961 |
| Validation | | | | | |
| MolProbity score | 1.82 | 2.02 | 1.92 | 2.30 | 2.16 |
| Clashscore | 7.0 | 7.13 | 6.25 | 8.19 | 7.85 |
| Rotamer outliers (%) | 1.9 | 2.6 | 2.3 | 4.2 | 3.5 |
| Ramachandran plot | | | | | |
| Favored (%) | 96.5 | 95.5 | 95.7 | 94.3 | 95.4 |
| Allowed (%) | 3.5 | 4.4 | 4.3 | 5.6 | 4.5 |
| Disallowed (%) | 0 | 0.1 | 0 | 0.1 | 0.1 |
